# Region-Level Design and Analysis of CRISPR Perturbation Screens with FRACTEL

**DOI:** 10.64898/2026.07.06.736794

**Authors:** Richard W Doty, Maria A. ter Weele, Alejandro Barrera, Lexi R. Bounds, Charles A. Gersbach, Andrew S. Allen

**Author notes:** These authors contributed equally to this work (Richard W Doty, Maria A. ter Weele, Alejandro Barrera).

## Abstract

We present FRACTEL, a statistical framework for region-level analysis of CRISPR perturbation screens. FRACTEL aggregates gRNA *p*-values using a bounded minimum across order statistics, preserving scale and enabling adaptive sensitivity to sparse or diffuse effects. Region-level null distributions are estimated via simulation, ensuring precise type I error control. Simulations and real CRISPRi/a datasets demonstrate improved power and replication rate over gRNA-level analyses. FRACTEL also informs experimental design, revealing trade-offs between gRNA redundancy and efficacy and identifying inherent limits in single-cell repression screens of lowly expressed genes. The method integrates with existing pipelines and supports diverse CRISPR screening applications.

## Introduction

High-throughput CRISPR screens have emerged as a scalable experimental approach for dissecting the effects of genomic perturbations on diverse phenotypes, including gene expression^1–6^, migration^7^, and cell growth^5,8–15^. Initial applications focused on CRISPR-Cas9 knockout screens in coding regions, producing insertions and deletions with variable efficiency, and disrupting gene function with incomplete penetrance and typically large effect sizes^10,16–18^. Expansions to CRISPR interference (CRISPRi) and activation (CRISPRa) screens with diverse epigenetic modifiers have facilitated targeting noncoding regulatory elements, reading out subsequent effects via cell growth or via gene expression of reporter constructs^1,19–21^. More recently, CRISPR screens coupled with single-cell RNA sequencing (scRNA-seq) have mapped regulatory elements to genes in particular loci or genome-wide^9,16,22–30^. These Perturb-seq screens have identified regions crucial for controlling expression of specific genes (often referred as regulatory elements), elucidated gene networks, and linked genetic elements to phenotypic traits and eQTLs^22–24,28^. While the noncoding genome makes up >98% of human DNA and contains ∼90% of trait-associated genetic variation, the regulatory logic of active elements remains poorly understood. Efforts like the ENCODE and Roadmap consortiums have mapped the activity of the noncoding genome and identified putative regulatory elements across cell types^31,32^; CRISPRi/a screens have become a standard approach for dissecting the effect of these putative regulatory elements in their native genomic and cellular contexts^33^.

A hallmark of CRISPR perturbation screens is the use of multiple guide RNAs (gRNAs) to target the same putative regulatory element. This redundancy increases the likelihood of successful perturbation despite variability in gRNA efficacy or specificity^27^, a persistent challenge in CRISPR-based assays and similarly addressed in short hairpin RNA (shRNA) or CRISPR knockout screens^5,13^. However, it also raises a statistical issue: most analyses are performed at the gRNA level, i.e. the analysis estimates the effect of a particular gRNA, not the element itself. While it might be tempting to assign significance to a region if at least one of the gRNAs targeting within the region is individually significant, this does not control type I error at the region level and can substantially reduce power when multiple gRNAs show moderate but consistent effects.

Several aggregated region-level analysis methods have been proposed to address these shortcomings and estimate the effects of entire regulatory elements, rather than individual gRNAs. MAGeCK’s^34^ robust rank aggregation (RRA) method has been widely applied in bulk CRISPR screens, ranking gRNAs across the experiment to score genes or regions. scMAGeCK^35^ extends this approach to single-cell data, while methods such as CASA^2^, RELICS^36^, and CRISPR-SURF^37^, use positional information to model local perturbation activity. Alternative strategies, such as union-based pseudobulked aggregation or Bonferroni correction across gRNAs within a region, have also been implemented in pipelines like SCEPTRE^38,39^. While valuable, these methods have important limitations: rank-based approaches discard scale information that is present in the magnitude of the statistics and lose sensitivity in focused screens; tiling models can be underpowered when few gRNAs target a region or when gRNA effects are not linearly correlated; naïve multiple-testing corrections can be overly conservative.

These constraints are especially pronounced in scRNA-seq Perturb-seq CRISPR screens, where experimental design is shaped by limited cell numbers, sequencing depth, and the high cost of coverage across large gRNA libraries. While having multiple gRNAs per element increases confidence, it multiplicatively increases the number of cells (and subsequent sequencing) necessary to maintain statistical power per perturbation. Sequencing costs, gRNA variability, and limitations in cell number have incentivized development of compact optimized CRISPR gRNA libraries^10,14,40–45^. However, novel noncoding regions without validated gRNAs still require 10-20 novel gRNAs per element^5^. As we demonstrate, trade-offs between gRNA redundancy and per-gRNA coverage can have profound effects on statistical power, particularly for modest effect sizes or lowly expressed genes. Furthermore, in repression assays targeting genes near the detection limit (e.g., lowly expressed genes), even strong perturbations can yield data that are difficult to analyze due to stochastic dropout and complete separation in count-based models.

To address these challenges, we developed FRACTEL (Framework for Rank Aggregation of CRISPR Tests within ELements), a region-level statistical framework that aggregates gRNA-level *p*-values using a bounded minimum across order statistics. FRACTEL preserves the scale of *p*-values, allows bound tuning to adapt to sparse or diffuse signal architectures, uses pre-computed null simulations, and controls element-level type I error. This approach integrates seamlessly with existing gRNA-level inference methods, offering a generalizable and computationally efficient approach to region-level inference. In addition to improving discovery power, FRACTEL provides a principled framework for quantifying the impact of experimental design choices—such as the number and efficacy of gRNAs—on statistical performance.

## Results

### Overview of the FRACTEL approach

FRACTEL is designed to improve region-level inference in CRISPR perturbation screens by aggregating statistical evidence across multiple gRNAs targeting the same genomic element (Fig. 1). For each element, *n* gRNAs are tested to obtain individual *p*-values. We assume that these *p*-values are generated from a statistical procedure, such as SCEPTRE, that produces well calibrated *p*-values so that they are uniformly distributed under the null. In cases where there is miscalibration, but where nontargeting guides are available, we give an adjustment procedure that empirically rescales the *p*-value distribution using the nontargeting guides (see Methods). We then order the *p*-values from smallest to largest. Under the null hypothesis, the *i*^th^ ordered *p*-value, *P*_(i)_, follows a Beta distribution parameterized by its rank, i.e.,

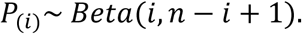

**Figure 1.**
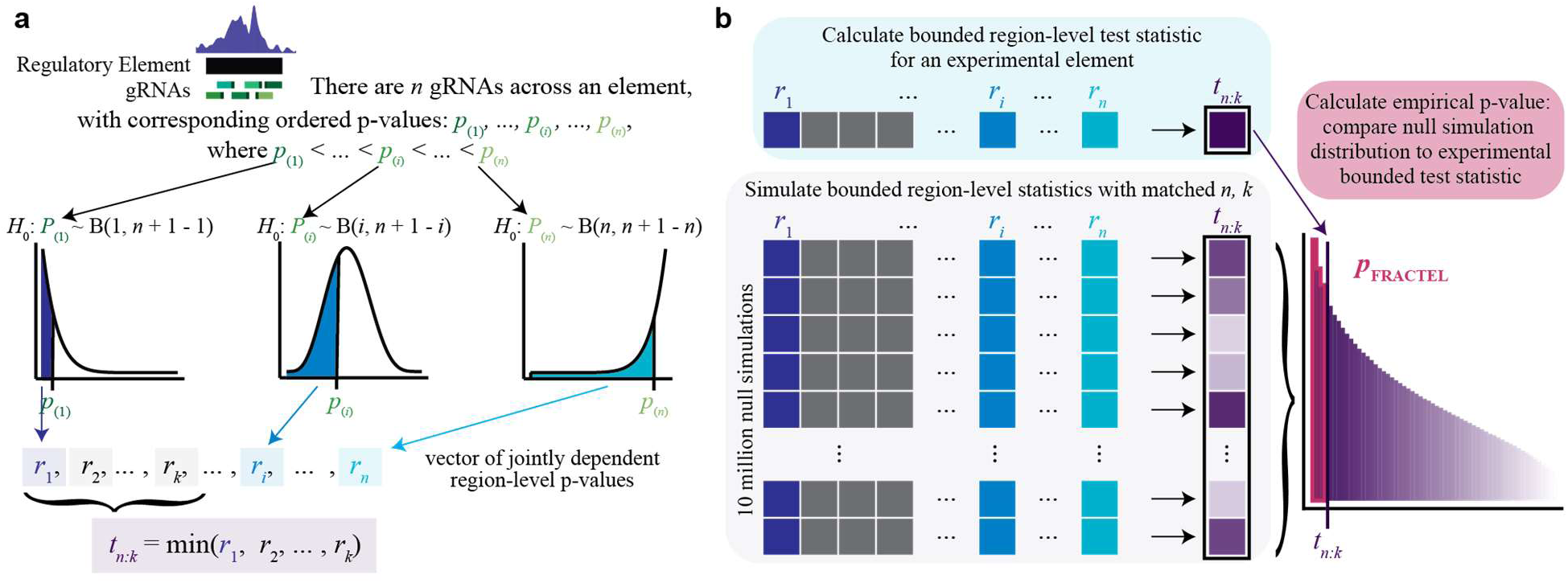
Overview of the FRACTEL framework for element-level inference in CRISPR screens. **(A)** For each regulatory element targeted by *m* gRNAs, individual gRNA-level *p*-values are obtained and ordered (*p*_(l)_ < ⋯ < *p*_(*n*)_). Under the null hypothesis, the *i*-th ordered *p*-value follows a *Beta*(*i*, *n* + 1 − *i*) distribution. Each ordered *p*-value is evaluated under its corresponding Beta model to yield a set of element-level *p*-values (*r*_l_, …, *r*_n_). The bounded-minimum statistic (*t_n:k_*) is then defined as the smallest value among the first *k* element-level *p*-values. **(B)** FRACTEL estimates the null distribution of this statistic through large-scale simulations matched to the observed number of gRNAs. The empirical *p*-value (p_FRACTEL_) is obtained by comparing the observed bounded statistic to 10 million simulated nulls, ensuring accurate type I error control across diverse effect architectures.

FRACTEL evaluates each *p*_(i)_ under this distribution to produce a new *p*-value, *r*_i_, that quantifies how unusual it is, under the null, for the *i*^th^ ordered *p*-value out of the set of *n* to be as small as *p*_(i)_. Note that *R*_i_∼ *Uniform*(0,1) under the null hypothesis, but the collection is jointly dependent. We denote these observed *p*-values as *r*_i_ to make it explicit that each of these is capturing deviations from the null at the region level. We then use a bounded minimum on (*r*_l_, …, *r*_k_) to denote the element-wise test statistic

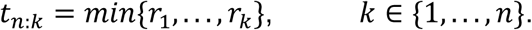

The bound parameter *k* controls the trade-off between sensitivity to concentrated (strong) and distributed (moderate) effects (Fig. 1a). Alternatively, *k* can be derived from a user-specified proportion *b* where *k* is defined as the smallest integer greater than or equal to *b×n*.

For each element, the FRACTEL *p*-value is computed by comparing the observed *t_n:k_* to the null distribution of the bounded-minimum statistic estimated via Monte Carlo simulation conditional on the number of gRNAs, *n*, enabling precise control of type I error without permutation of the observed data (Fig. 1b). FRACTEL is compatible with any gRNA-level testing framework that yields calibrated *p*-values, allowing seamless integration into existing analysis pipelines.

In the following sections, we evaluate FRACTEL’s calibration, power, and biological performance using first simulated data and then real CRISPRi/a Perturb-seq screens. We also explore how experimental design parameters — including the number of gRNAs, gRNA efficacy, and target gene expression level — influence detection power, and we compare FRACTEL to SCEPTRE, a widely-used powerful and FDR-controlled method for Perturb-seq analysis^38,39,46^.

### Control of type I error under the regional null

We evaluated FRACTEL’s type I error control under the regional null hypothesis that all gRNA-level effects for a given region are null. We simulated this situation by independently drawing individual gRNA *z*-scores for a given region from a standard normal distribution, yielding uniform *p*-values. For each element, we computed FRACTEL *p*-values for varying numbers of gRNAs per element *n* and bound parameter *k* where *k* is either a specified integer or *ceiling*(*b* ∗ *n*) where *b* is a user-specified proportion between 0 and 1.

Null distributions were generated from 100,000 simulated replicates for each *n* and repeated across baseline-expression settings. In our model, baseline expression corresponds to the mean count when *x*_ij_ = 0, given by *l̄e*^/30^ (see Methods). We fixed the mean library size at *l̄* = 300 and varied *β*_0_ ∈ {-6,-4,-2}, corresponding to mean UMI counts of 0.744, 5.495, and 40.601, respectively. Empirical *p*-values were calculated as the proportion of simulated alternatives below α = 0.05; correct calibration was expected to yield rates near this threshold. Across all tested (*n*, β₀) combinations (*n* = 5–20, *β*_0_ ∈ {-6,- 4,-2}), observed type I error rates closely matched the nominal level. For example, the empirical error rate was 0.0501 with *n* = 5 and high baseline expression, and 0.0494 with *n* = 20 and low baseline expression (Table S1).

### Impact of bound parameter choice

The bound parameter *k* in FRACTEL determines how many of the smallest ordered gRNA-level *p*-values contribute to the bounded-minimum statistic. Smaller values of *k* favor detection of sparse, strong effects concentrated in a few gRNAs, whereas larger values allow moderate effects to accumulate across many gRNAs, improving sensitivity for diffuse signal architectures.

We used simulations to examine how power varies with *k* across different underlying configurations of active gRNAs. For each scenario, we fixed the total number of gRNAs (*n* = 10) and varied the number of active gRNAs, *q*, from one to ten. Power increases with the number of active gRNAs as expected, and the *k* value that maximizes power shifts upward accordingly (Fig. 2a). When signal is sparse, optimal power is achieved with small *k* (*k* ≈ 0.2*n–0.3n*), whereas more distributed effects favor larger *k* (*k* ≈ 0.6*n*–0.8*n*).

**Figure 2.**
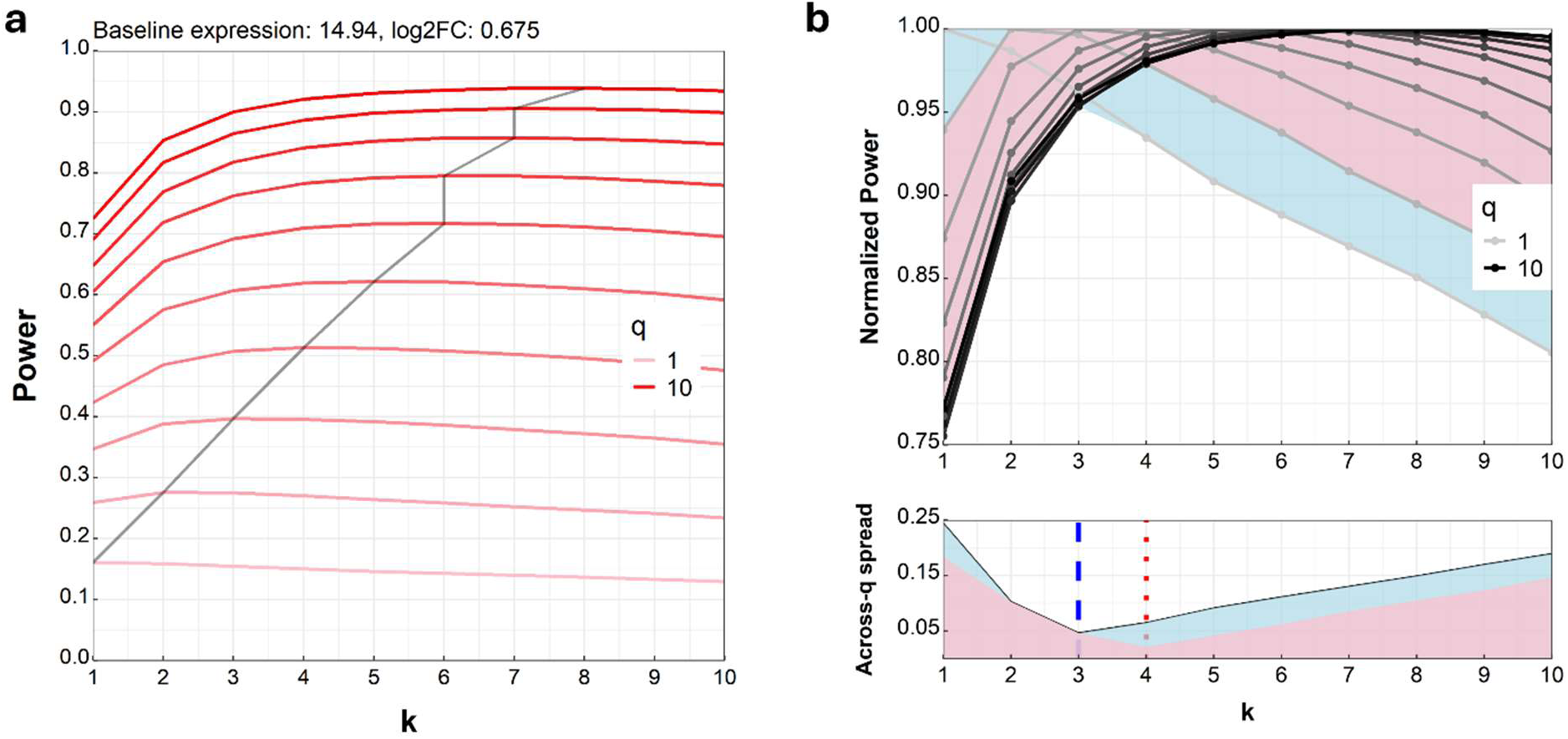
Effect of the bound parameter on power across signal architectures. **(a)** Power as a function of the bound parameter *k* for elements with different numbers of working gRNAs (*n* = 10). Each line represents a distinct number of active gRNAs, from one (red) to ten (light pink). The black line traces the maximum power achieved for each configuration, showing that the optimal *k* increases with the number of working gRNAs. **(b)** Normalized power curves corresponding to panel (a), where each line is scaled by its maximum power across *k*. Each curve therefore reflects relative performance as *k* varies for a fixed number of active gRNAs. The shaded region represents the range of normalized power across all architectures at each *k*. The lower subplot shows the width of this range (max–min) as a function of *k*. In this setting, the spread is minimized near *k* = 3 (blue dashed line), indicating stable near-optimal performance across the simulated signal architectures. The red dashed line marks *k* = 4, which may be preferable when there is prior expectation that more than one gRNA contributes to the signal.

Although the optimal *k* depends on the true signal architecture (the number of gRNAs modulating the regulatory element), *k =* 0.3*n–*0.4*n* provides a practical robust trade-off when that structure is unknown. Rather than selecting *k* to maximize power for any single assumed architecture, we use this range to reduce sensitivity to misspecification of the number of working gRNAs. In the example simulation used for Fig. 2, the narrowest spread in normalized power occurs near *k* = 3, indicating consistent near-optimal performance across architectures. This value therefore balances sensitivity and stability, serving as a practical default for most CRISPRi/a screen designs. Importantly, the optimal choice of *k* depends on the design architecture and prior expectations (Fig. S1). For example, if there is a strong prior belief that more than one gRNA is effective, the optimal choice for *k* becomes *k* = 4 (Fig 2b).

### Power under varying gRNA number and efficacy

While many compact validated genome-wide promoter-targeting libraries have been generated^10,14,40–45^, noncoding screens still require 10-20 novel gRNAs per element and typically have much smaller effect sizes on gene expression^5^. Screening with more gRNAs per regulatory element can either sharpen or blur the signal, depending on how many of those gRNAs are genuinely functional. To examine this trade-off, we compared designs with ten and twenty gRNAs per element under identical baseline parameters, varying the number or proportion of working gRNAs while sweeping effect sizes across a broad fold-change range (in log scale). For each configuration, we estimated power using the FRACTEL test and then computed pointwise differences between the *n* = 10 and *n* = 20 curves where positive values indicate higher power for *n* = 10.

When the number of effective gRNAs was held constant, the designs with a smaller number of total gRNAs consistently maintained higher power to detect non-null effect sizes (Fig. 3a). The positive power difference between designs with *n* = 10 and *n* = 20 reflects dilution by non-functional gRNAs in the larger library. As the number of working gRNAs increased, this advantage became progressively narrower—the peaks in the blue curves contract—because stronger cumulative signal allowed both designs to reach power saturation (detection probability approaching one) at ever-smaller effect sizes. In parallel, the *n* = 10 design becomes functionally saturated once all gRNAs are active, while *n* = 20 still contains half inactive gRNAs. Even near power saturation, *n* = 10 retains a slight edge due to its cleaner signal composition. The central dip at log_2_FC = 0 corresponds to both designs returning to baseline detection rates near 0.05, as expected for a well-calibrated 0.05-level test.

**Figure 3.**
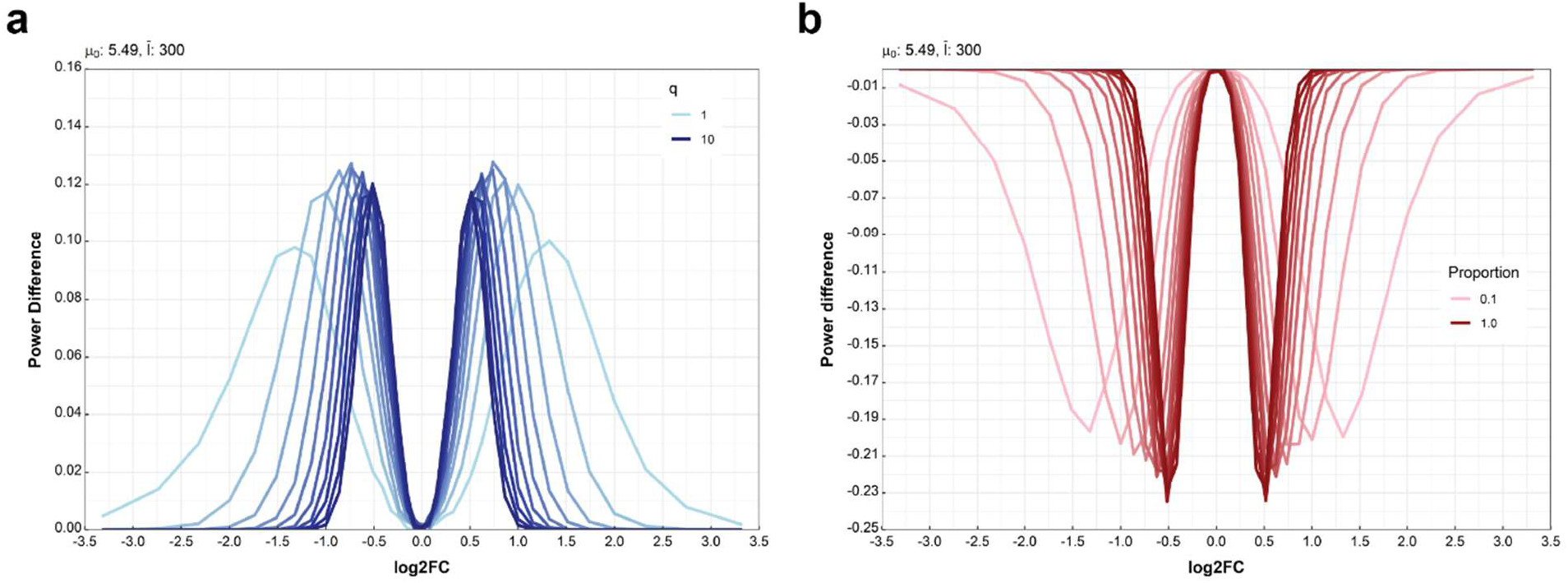
Effect of gRNA number on power under fixed numbers and proportions of functional gRNAs. **(a)** Power difference (*ΔP* = *P*_n=l0_ − *P*_n=20_) across log2 fold-change effect sizes for designs with ten versus twenty gRNAs per element, holding the number of working gRNAs fixed (*q* = 1–10). Each blue curve represents a distinct value of *q*. Positive values indicate higher power for *n* = 10, reflecting dilution by non-functional gRNAs in the larger library. As *q* increases, the peaks contract, indicating that the advantage of the smaller design diminishes as both configurations approach power saturation. The central dip near log2FC = 0 corresponds to both designs returning to baseline detection rates near 0.05. **(b)** Power difference under fixed proportions of working gRNAs (*q*/*n*). Each curve represents a distinct working-guide proportion. Negative values indicate higher power for *n* = 20. When the proportion of functional guides is held constant, increasing the total number of guides increases the absolute number of active perturbations, leading to higher power for the larger design. Together, panels (a) and (b) show that increasing guide number improves detection only when it increases the number of functional gRNAs.

When the proportion of working gRNAs was fixed instead (Fig. 3b), the larger library outperformed the smaller one. In this setting, *n* = 20 includes more effective gRNAs in absolute terms, and the gain in signal outweighs the added noise, leading to consistently higher power across effect sizes. Taken together, the two panels illustrate the same underlying trade-off from complementary perspectives: increasing the number of gRNAs improves detection only when it increases the number of functional perturbations. Expanding coverage across an element can raise the likelihood of including effective guides, but the benefit diminishes if additional guides are increasingly likely to be inactive, as the added noise offsets the expected gains.

### Constraints from basal gene expression in repression screens

Detection of strong repression at very low basal gene expression levels faces two distinct constraints—one practical, the other theoretical—that together define the lower boundary of statistical power for CRISPRi analyses based on maximum-likelihood estimation.

The practical limit arises when transcript abundance in the perturbed group is so low that few or no molecules are detected. In these cases, model fitting becomes unstable or fails altogether due to complete or near-complete separation of counts between conditions. In effect, when all measured counts in the perturbed (repressed) group fall to zero, the experiment ceases to provide the information required for statistical inference, regardless of the statistical method used. Analysis workflows often exclude such element-gene pairs during quality control or testing^38,47^, which leads to a systematic bias against detecting strong repressive effects.

The theoretical limit persists even when detection is idealized. We conducted multiple simulations in parallel to assess the effect of baseline expression on power in the ideal case where test statistics were generated directly, removing measurement errors (Fig. S2). The simulations directly show clear asymmetry for small values of *β*_0_ (baseline-expression) with the difference between activation and repression disappearing as *β*_0_ increases. The underlying mechanism is inherent to maximum-likelihood estimation. As the true fold-change approaches zero, the estimated variance grows much faster than the change in mean, forcing the expected test statistic toward zero. This divergence is evident in the simulated curves, where the standard error inflates exponentially (Fig. S3). The system effectively enters a quasi-Bernoulli regime in which detection depends on whether any perturbed cell retains measurable transcript counts.

Together, these findings delineate the limits of inference for low-expression targets. The practical constraint restricts what can be measured, and the theoretical constraint limits what can be recovered even under perfect measurement.

### Targeted vs transcriptome-wide designs

Single-cell CRISPR screens can be performed using either transcriptome-wide profiling or targeted enrichment panels that capture a defined subset of genes^9,48^. Transcriptome-wide designs support unbiased discovery but distribute sequencing reads across thousands of transcripts, limiting sequencing coverage for individual targets. Targeted approaches concentrate reads on selected genes, improving the effective detectability of perturbation effects.

In our modeling framework, enrichment operates through the mean library size term *l̄*, which scales the expected transcript count for each cell. Because the model described in Methods defines expression as

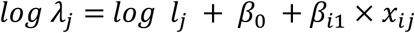

an increase in *l̄* has the same multiplicative effect on mean expression as an increase in the baseline expression parameter *β*_0_. Mathematically, the two are indistinguishable—both shift the expected count distribution upward and thereby move genes into a higher-powered regime.

This equivalence allows targeted enrichment to be understood as a practical mechanism for increasing the effective baseline expression of genes within a panel. Simulations varying *β*_0_ thus capture the same statistical consequences as changes in sequencing depth or panel coverage. Increasing *β*_0_ (or equivalently, *l̄*) shifts power curves inward, indicating greater power for detecting smaller fold-changes at greater baseline expressions (Fig. S2). Power gains are most pronounced for genes near the detection limit under standard sequencing depth and diminish for highly expressed genes that are well-powered even without enrichment.

However, enrichment designs introduce new considerations. Like transcriptome-wide sequencing, targeted panels impose competitive expression, but even more noticeable: a gene’s apparent abundance is measured relative to other panel genes, not total transcriptome output. This means that adding highly expressed genes to a panel can dilute signal from lower-abundance targets. From a modeling perspective, this compresses the effective dynamic range and complicates the interpretation of *β*_0_ across datasets with different panels.

These considerations frame targeted enrichment not as a separate statistical scenario but as an experimental route to the same outcome achieved by higher baseline expression—enhanced sensitivity for lowly and moderately expressed genes when sequencing resources are efficiently allocated.

### Benchmarking on real datasets

FRACTEL can be applied to aggregate any well-calibrated gRNA-level tests to produce element-level analyses of CRISPR screens. Many technical factors and model misspecification can confound CRISPR screen analysis, producing poorly calibrated analyses with high numbers of false positives across common analysis methods and foundational single-cell CRISPR screen datasets^39^. However, the conditional resampling methods proposed by Barry and colleagues (SCEPTRE) robustly address miscalibration in real-world data, producing well-calibrated *p*-values for individual gRNA-gene pairs across datasets. These are appropriate inputs for FRACTEL.

We applied FRACTEL to SCEPTRE gRNA-gene pair results from five CRISPRi/a single-cell datasets targeting a union set of putative noncoding regulatory elements in K562s, induced pluripotent stem cells (iPSCs) and neural progenitor cells (NPCs) (see Methods, **Supplementary Tables X1-X5**)^48^. These screens targeted the major histocompatibility complex (MHC) locus with the same library of 20 gRNAs per putative regulatory element, various cell-type specific positive control promoter regions targeted by 10 gRNAs each, and non-targeting gRNAs as negative controls. As these gRNAs were untested in previous assays, variability between gRNAs targeting the same element was expected. SCEPTRE provides two options for region-level analysis in their computational methods: a Bonferroni correction between gRNAs appropriate for regions with high variability between gRNA effects, and a “union test” approach that pseudobulks gRNAs targeting a region together, powerful for cases with little variability between gRNA-gene effects across a region. We used both these methods as benchmarks for comparison with FRACTEL’s aggregation. Simulation results illustrate the limitations of the union test under increasing heterogeneity in gRNA effects, where opposing or uneven contributions can attenuate aggregate signal (**Fig. S4**). We also see that FRACTEL performs no worse than the Bonferroni approach across experimental designs (**Fig. S5**). We propose FRACTEL as a strategy with more flexibility than the union test or Bonferroni correction, robust to some variance between gRNAs but not entirely dependent on the best gRNA per element.

Positive control pairs were robustly detected by all three approaches across all screens (**Fig. 4a**, **Fig. S6a, Supplementary Tables X6-X10**). In CRISPRi screens at small FDRs (FDR < 0.01), the union approach made slightly more discoveries (**Fig. 4a-b**, **Fig. S6a-b**). At larger FDRs (FDR = 0.05 to 0.1), FRACTEL detected more significant element-gene pairs than either a Bonferroni correction or union approach, across all datasets (**Fig. 4b**, **Fig. S6a-d(ii)**). As an example, the CRISPRi screen of the MHC locus in iPSCs had 304 FRACTEL discoveries, with 69% shared by at least one other method, while the union approach had 269 discoveries, with 64% shared by another method (FDR = 0.05, **Fig. 4**). As expected, element-gene pairs uniquely identified by FRACTEL or the Bonferroni correction had considerably more variable effect sizes across gRNA-gene pairs within the element-gene pair than the union approach (**Fig. 4c**, **Fig. S6a-d(iii)**). While this is an advantage for detecting functional genomic elements with sparse gRNA signal, FRACTEL, like a Bonferroni correction, is sensitive to off-target or unusual effects driven by one highly significant gRNA.

**Figure 4.**
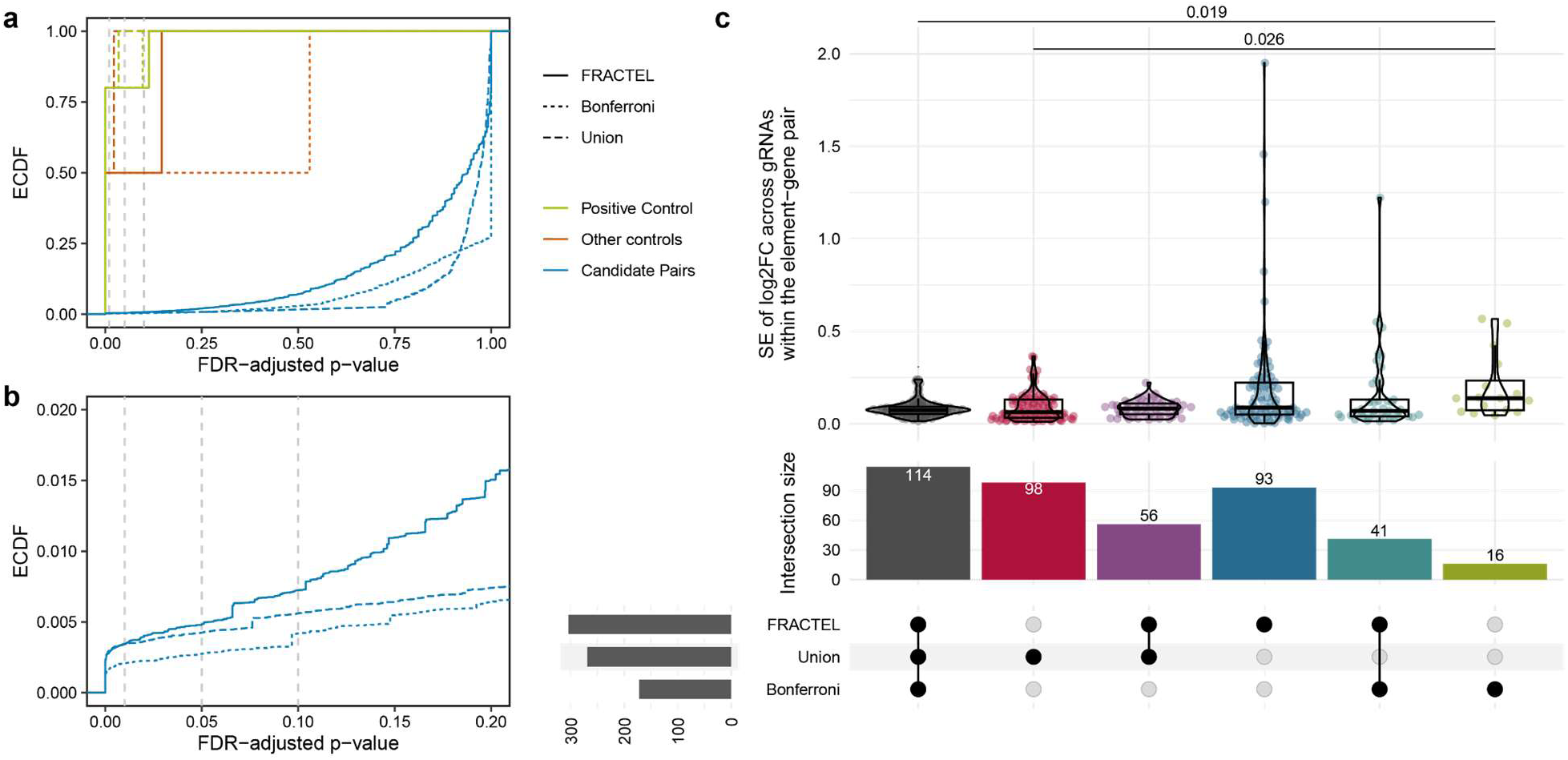
Comparison of aggregation methods. **(a)** Empirical cumulative distribution function (ECDF) of the three aggregation methods across FDRs in a dCas9-KRAB Perturb-seq screen of the MHC locus in iPSCs^48^. **(b)** Plot of ECDF in **(a)** for non-control elements with FDR < 0.2. Dashed vertical grey lines in **(a-b)** represent FDRs of 0.01, 0.05, and 0.1. **(c)** Comparison of standard error of the effect sizes (*log*_2_ (*FC*)) between gRNAs within an element-gene pair, for exclusive intersection sets of significant (FDR < 0.05) elements called by each of the three methods, along with an upset plot to show set sizes. Significant pairwise Mann-Whitney U-tests with a Holm correction are displayed (*p*_adj_ < 0.05). Not shown is the exclusive intersection of the union and Bonferroni methods (*n* = 1).

To further understand differences between the three approaches and their sensitivity to individual gRNAs, we plotted the distributions of ordered *p*-values (*p*_(i)_) from SCEPTRE’s well-calibrated individual gRNA-gene tests, separated by which tests detected the element-gene pair as significant, alongside the set of insignificant *p*-values for comparison (**Fig. S7**). As expected, the Bonferroni correction is sensitive to only the most significant gRNA in an element: element-gene pairs detected by a Bonferroni test have a very small *p*_(l)_, but with much larger *p*_(2)_ and *p*_(3)_ than any of the other significant elements. Element-gene pairs called exclusively by the union approach had a larger value of *p*_(l)_, consistent with the union approach being less dependent on the singular best gRNA. Counterintuitively, these pairs called exclusively by the union method also tended to have larger values of *p*_(2)_ and *p*_(3)_: Though we expected these elements to have more consistently significant gRNA effects, gRNAs do not have to be individually very significant for pseudobulked methods.

To examine these further, we plotted element-gene pairs called only by the union test or FRACTEL (**Fig. S8, S9**). Element-gene pairs called by only the union test (FDR_Union_ < 0.05, FDR_FRACTEL_ > 0.2) tended to have less significant gRNAs with small effect sizes that all trend in the same direction (**Fig. S8**). Some element-gene pairs called highly significant by the union approach were not recovered by FRACTEL until a much larger FDR; these fell into 2 groups. The first group contained element-gene pairs with a very small number of cells per gRNA-gene pair, driven either by low gRNA coverage or low target gene expression: the top gRNA-gene tests in these element-gene pairs have little power and thus lack sufficient significance to be detected with FRACTEL. The second group contained element-gene pairs with very mild but consistent gRNA-gene effects across the whole element, making the element-gene pair significant without any singularly significant gRNA. This second group of element-gene pairs and the difference between the union and FRACTEL result is reduced by raising the bound parameter so that more gRNAs are considered within the FRACTEL test (**Fig. S10, Additional Files 11-15**).

Element-gene pairs called only by FRACTEL (FDR_FRACTEL_ < 0.05, FDR_Union_ > 0.2) also fell into 2 groups, both of which necessarily had a sufficiently significant gRNA within the tunable bound. The first was element-gene pairs in which one gRNAs’ effect dominates the entire element (e.g. *CLIC1*-chr6:31,111,005-31,111,190, **Fig. S9**). The second group had significant gRNAs with opposing directions of effect (e.g. *EGFL8*-chr6:33,136,422-33,136,837, **Fig. S9**). Using two-sided SCEPTRE *p*-values as input to the aggregation makes FRACTEL sensitive to elements with gRNAs that have significant but opposing directions of effect, which a union approach would always miss.

## Discussion

CRISPR-based perturbation screens have become indispensable tools for mapping the regulatory architecture of the genome, yet most analytical pipelines have been optimized to perform gRNA-level inference. This focus overlooks the fact that the primary unit of biological interest is the genomic region, leading to inflated region-level false discovery rates and reduced power when signals are distributed across multiple gRNAs. Furthermore, these negative effects can be amplified in increasingly common downstream analyses focused on the association of perturbations with identified cellular programs and networks. Our study introduces FRACTEL, a statistical framework that addresses this disconnect by aggregating gRNA-level evidence into principled region-level inference.

Our approach offers several key advantages and some limitations. FRACTEL preserves the scale of gRNA-level *p*-values rather than normalizing them, allowing more information to be retained during aggregation while maintaining the FDR. In addition, robust rank aggregation is agnostic to the input test, allowing for future improvements in gRNA-level differential expression analysis that further address biological heterogeneity or technical factors. However, FRACTEL *requires* well-calibrated gRNA-level *p*-values, and many differential expression analyses inflate false-positives^39^, making them inappropriate inputs to FRACTEL. As such, we recommend using gRNA-level analysis from SCEPTRE as input to FRACTEL for scRNA-seq CRISPR screens, as those are typically well-calibrated. Alternatively, we provide an empirical *p*-value transformation to enable calibration using negative controls (see Methods). Although we focus on statistical significance in our analyses here, we also provide the computation of a weighted average effect estimate in our code (see Methods)

FRACTEL is sensitive to a wide range of gRNA effect architectures by tuning the bound of aggregation, favoring either sparse, strong effects or diffuse, moderate effects depending on the underlying biology. Our recommendation of a bound of *k* = 0.3*n* or 0.4*n* is a balance between power and expected sparsity: we assume not all gRNAs have the same efficacy, an assumption appropriate for contexts where gRNA potency is less predictable (e.g. enhancer screens with previously untested gRNAs), but less necessary for screens with only high efficacy gRNAs (e.g. promoter libraries of validated or pre-screened gRNAs). As the field evolves to more confident predictions of gRNA specificity and strength, models should adapt to explicitly account for those expectations; for FRACTEL, that might mean an adaptive bound per element rather than a singular experimental bound. We show by simulation that, with little variation between gRNAs, a union approach will remain more powerful than FRACTEL with a low bound (*k* = 0.3*n*) for small or moderate effect sizes. However, this difference is reduced by increasing FRACTEL’s bound.

FRACTEL constructs empirical null distributions using large-scale simulation, which can be precomputed for each gRNA count and reused across experiments. This strategy substantially reduces computational overhead while maintaining precise control over false positive rates. While practical, the use of a simulated null creates empirical *p*-values with limited precision. Future work should address this and allow analytical computation of exact *p*-values to improve reproducibility in the tail.

Several practical considerations emerged from our analyses. Repression screens targeting genes with very low baseline expression face an inherent ceiling on statistical power. As repression drives expression toward zero, complete separation inflates standard errors, reducing significance even for large effect sizes. This limitation reflects the biology of sparse transcript detection rather than shortcomings of the model, underscoring the importance of pre-filtering low-expression targets or incorporating targeted enrichment strategies. FRACTEL also assumes approximate independence of gRNA-level tests, which is reasonable for most CRISPRi/a designs but may be violated in cases of overlapping gRNAs or shared sequence features; in these cases, other models designed for tiling experiments (e.g. CRISPR-SURF) may be more appropriate^37^.

By enabling power analysis, our results also highlight design trade-offs between gRNA redundancy and efficacy. Adding non-functional gRNAs introduces noise and reduces power, while increasing the number of effective gRNAs substantially improves detection. These findings argue for broad coverage in exploratory screens to maximize the chance of including effective gRNAs, followed by validation experiments that prioritize high-efficacy gRNAs. Incorporating predictive models of gRNA activity into library design may further optimize this balance.

Here we have presented FRACTEL as a means to aggregate repeated tests of single effects (expression of a single gene). While differential expression testing is a core analysis for scRNA-seq CRISPR screens, it considers only one gene at a time, not broader transcriptional profiles. For perturbations with broad effects on cell states, like transcription factor perturbations, analyses that consider more than one transcriptional output at a time (e.g. energy distance clustering and embedding) may be more powerful than examining only one gene at a time^28,49^. FRACTEL could also be used to aggregate across multiple elements or genes to create gene program-level scores over pre-determined gene sets.

Although developed for CRISPR screens, FRACTEL may be relevant more broadly. Rank-based aggregation of calibrated test statistics could support inference in other high-dimensional contexts, such as enhancer screens, eQTL hotspots, or spatial transcriptomics. By providing both a methodological advance and a framework for design analysis, FRACTEL contributes to improving the robustness and interpretability of perturbation-based genomics.

## Conclusion

We developed FRACTEL, a statistical framework for element-level inference in CRISPR perturbation screens. By aggregating gRNA-level *p*-values through a bounded-minimum approach and calibrating the null distribution via simulation, FRACTEL achieves accurate type I error control while retaining sensitivity across diverse effect architectures.

Our simulations and applications to real CRISPRi/a datasets demonstrate that FRACTEL improves discovery power compared to existing methods, while remaining tunable to different effect architectures. In addition, FRACTEL provides actionable guidance for experimental design, clarifying how gRNA number, efficacy, and target gene expression influence detection power. These insights highlight both the opportunities and intrinsic limitations of CRISPR-based repression assays, particularly for lowly expressed genes.

As a modular and computationally efficient method, FRACTEL integrates seamlessly with existing gRNA-level testing frameworks. Its adoption will enhance reproducibility and interpretability in CRISPR perturbation studies, while also providing a foundation for broader applications in high-dimensional genomics where multiple statistical tests must be combined to capture region-level effects.

## Methods

### FRACTEL test setup

We aim to test whether a regulatory element is associated with a response by aggregating evidence across its gRNAs. Suppose an element is targeted by *n* gRNAs. For a given element in a single-cell assay let *Z*_l_, …, *Z*_n_ denote the gRNA-level test statistics and *p*_l_, …, *p*_n_ their corresponding *p*-values. We assume only that the gRNA-level *p*-values are valid under the null hypothesis; we do not otherwise constrain how they are obtained. Let *p*_(l)_ ≤⋅⋅⋅≤ *p*_(*n*)_ denote the ordered *p*-values.

Calling a region based solely on the minimum *p*-value, *p*_(l)_, is powerful when signal is extremely sparse but can lose power when effects are modest and shared across multiple gRNAs. Our goal is to aggregate ordered evidence in a manner that adapts to unknown effect architectures across gRNAs.

### Order-statistic transformation to region-level *p*-values

Under the regional null hypothesis, all *n* gRNAs have no effect and the *p*-values, *p*_l_, …, *p*_n_ are i.i.d. Uniform(0,1). Let *p*_(l)_ ≤⋅⋅⋅≤ *p*_(*n*)_ denote the ordered *p*-values. It is well known that the *i*^th^ order statistic satisfies

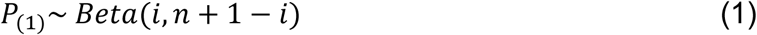

Applying the probability integral transform to each order statistic, we define

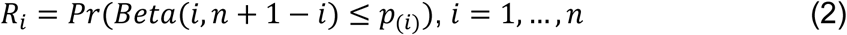

Note that *R*_i_ is marginally Uniform(0,1) under the null hypothesis, though the collection (*R*_l_, …, *R*_n_) is jointly dependent through the ordering of the original *p*-values.

Each *R*_i_ yields a legitimate element-level test attuned to different effect architecture (e.g., *i* = 1 favors sparse, strong signals; larger *i* favor modest effects shared across multiple gRNAs).

### Architecture-adaptive aggregation via a bounded minimum

To adapt across architectures without overcommitting to a single order statistic, we define a bounded minimum over the first *k* transformed *p*-values:

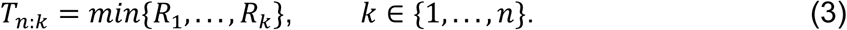

Smaller values of *k* emphasize sensitivity to sparse, large effects, while larger values increase power against more diffuse signal. We study the choice of *k* empirically and by simulation (see Simulations). The value of *k* used in the presented results is discussed in Impact of bound parameter choice.

### Null calibration and computation

Although each *R*_i_ is marginally Uniform(0,1) under the regional null hypothesis, the vector (*R*_l_, …, *R*_k_) is jointly dependent due to the ordering of the underlying *p*-values. As a result, the null distribution of *T_n:k_* does not admit a simple closed form. Importantly, under the regional null it depends only on the pair (*n*, *k*). We therefore approximate the null distribution of *T_n:k_* via Monte Carlo simulation for each observed *n* and chosen *k*, as follows:

1. Draw *n* i.i.d. Uniform(0,1) values and sort them to obtain *p*_(l)_, …, *p*_(*n*)_.
2. Compute *r*_l_, …, *r*_n_ via (2) and then *t_n:k_* via (3).
3. Repeat steps 1-2 a total of *N* times to form the empirical null distribution *F̂_n:k_*. The region-level *p*-value is then defined as

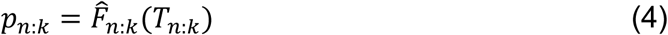

The assumption of independence among gRNA-level *p*-values is reasonable when the total gRNA library size is large relative to *n* and the experiment operates at low multiplicity of infection (MOI). Under these conditions, the probability that multiple gRNAs are present and active within the same cell is negligible, limiting induced dependence across gRNAs. If substantial dependence among gRNA-level *p*-values is anticipated, calibration via permutation or resampling may be substituted.

### Correcting Null Calibration

FRACTEL requires uniformly distributed negative control (non-targeting or safe-targeting) gRNA-level *p*-values. However, methods often produce many false positives^38,39^, with the extent varying by screen and analysis. We provide a computational tool to transform all *p*-values using a linear interpolation of the empirical cumulative distribution function (ECDF) of the non-targeting control gRNAs. This transformation measures the significance of each test gRNA as a linear interpolation of the proportion of non-targeting control gRNAs with smaller *p*-values than the test gRNA. This forces the background control *p*-value distribution to be uniform and reduces the significance of all other gRNAs, effectively measuring signal in the screen relative to internal negative controls.

### Effect size computation

Robust rank aggregation techniques such as FRACTEL provide p-values derived from aggregated test statistics. The thresholded FRACTEL statistic (*T_n:k_*) is therefore not itself a directly interpretable effect size, as it is also derived from element-level p-values. While one could compute the mean or median perturbation effect across the gRNAs within an element-phenotype pair, FRACTEL is explicitly designed to preserve power across both sparse and dense gRNA effect architectures, which can produce substantially different average effect sizes.

Instead, we consider each element as having either a predominantly positive or negative effect on the phenotype and compute two directional weighted averages of log_2_ fold changes using one-sided p-values, *p*^-^ and *p*^+^, as weights, taking the estimate with the largest absolute magnitude as the element-level effect size:

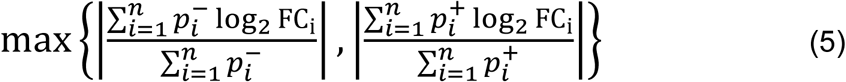

Using one-sided p-values as weights allows all gRNAs to contribute to the aggregate effect estimate according to both the magnitude and direction of their association with the phenotype. Because the reported effect size is defined as the directional estimate with the largest absolute magnitude, the resulting effect size is discontinuous around zero.

### Simulation studies

We evaluated the operating characteristics of the regional test *T_n:k_* across effect architectures and experimental designs.

### Generic gRNA-level simulation model

In order to assess the power of *T_n:k_* under a range of effect architectures, we simulate signal in a subset of gRNAs while keeping the remainder null. Let *q* ∈ {1, …, *n*} denote the number of non-null gRNAs (sparsity). We have noncentrality parameters *μ*_l_, …, *μ*_q_for the non-null gRNAs and set *μ*_q+l_ =⋅⋅⋅= *μ*_n_ = 0. We generate gRNA-level test statistics according to

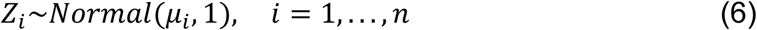

and compute one-sided *p*-values *p*_i_ = 1 − *Φ*(*Z*_i_). Region-level transformed *p*-values *R*_i_ and the bounded minimum statistic *t_n:k_* are obtained via (2)–(3). The resulting region-level *p*-value is obtained via (4).

We evaluate power over a grid of architectures defined by varying *q* and *n*. For non-null gRNAs, we consider homogenous effects (*μ*_l_=⋅⋅⋅= *μ*_q_) and heterogeneous effects, where *μ*_l_, …, *μ*_q_ are independently drawn from a *Normal*(*μ̅*, *τ*) distribution. The parameters *μ̅* and *τ*^2^ are explored over a grid.

### Linking *μ_i_* to experimental quantities (NB GLM → Z-scale)

To connect the noncentrality parameters *μ*_i_ used in the simulation model to interpretable CRISPR screen quantities, we consider a standard negative binomial generalized linear model for gRNA-level expression counts. Let *Y*_ij_ denote the UMI count for the target gene for gRNA *i* in cell *j*, let *x*_ij_ ∈ {0,1} indicate whether cell *j* carries gRNA *i*, and let *l*_j_ denote the library size (exposure). We model

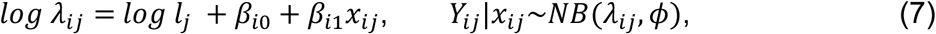

where *φ* ≥ 0 denotes the overdispersion parameter. The per-gRNA effect on the count scale is *FC*_i_ = *e^βil^*, with *FC*_i_ = 1 under the null. Note that *β*_il_ is on the natural log scale, not log_2_(FC).

Let *n*_il_ and *n*_i0_ denote the numbers of cells with and without gRNA *i*, respectively, and let *l̄* approximate the mean exposure (treating *log l*_j_ as an offset). Under standard regularity conditions, the Wald statistic *β̂*_*i*1_/*SE*(*β̂*_*i*1_) is approximately normal with unit variance. Its mean, which serves as the noncentrality parameter in (7) is therefore

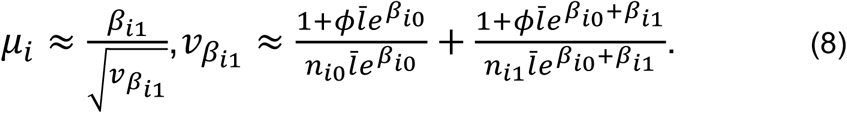

Equation (8) provides a direct mapping from experimental design parameters (*n*_*i*1_, *n*_*i*0_, *l̄*, *φ*) and effect size *β_i1_* = *log FC*_i_ to the Z-scale noncentrality *μ*_i_, which is then used as input to the simulation model in (6). The variance approximation *v_βi1_* follows standard results for negative binomial GLMs (see Zhu & Lakkis 2013^50^).

### Benchmarking studies

#### Datasets

Five CRISPRi/a screen Perturb-seq datasets targeting the MHC locus across 3 cell types (K562 cells, neural progenitor cells, and induced pluripotent stem cells) and 2 CRISPR effectors (dCas9-p300 and dCas9-KRAB) were obtained ^48^. The same gRNA library was used across all 5 screens, containing gRNAs targeting cell-type specific promoter controls (∼10 gRNAs per gene control), gRNAs targeting distal genomic putative regulatory elements (∼20 gRNAs per element), and 10% nontargeting gRNAs, for a final total of 12,723 gRNAs. These screens were conducted with probe-based targeted enrichment (see Bounds et al.^48^), enriching for transcripts in the MHC locus and some control genes outside the locus.

#### SCEPTRE Analysis

SCEPTRE was run on gene expression raw counts data generated by 10X Genomics CellRanger, as previously described^48^. Individual gRNAs were assigned to cells using CLEANSER^51^, a mixture model that estimates posterior probabilities for the presence of a gRNA in each gRNA-cell pair. Pairs with posterior probabilities over 0.5 were assigned, as in a previous analysis of this data^48^. SCEPTRE discovery pairs were constructed from a list of genes in the MHC locus and positive control genes (**Additional File 16**). In each case, negative control pairs were constructed from nontargeting gRNAs; notably, SCEPTRE constructs the negative control pairs from random groupings of nontargeting gRNAs in the Union and Bonferroni cases. Calibration checks for the five screens across the three SCEPTRE methods (Union, Bonferroni, and singleton analysis) are in Figure S10. Positive control pairs were not constructed separately from the discovery analysis and SCEPTRE’s native power check was not run; positive control pairs were annotated after running the discovery analysis and are all clearly detected (**Fig. 4a**, **Fig. S5**).

SCEPTRE was run with default covariates (`grna_n_nonzero, grna_n_umis, response_n_nonzero, response_n_umis`) in the high MOI setting (using the complement set of cells as the control group) with conditional resampling to produce 2-sided gRNA-gene *p*-values. SCEPTRE was run 3 times: once to produce gRNA-gene *p*-values and effect sizes, and twice more to produce SCEPTRE’s aggregated statistics for both the Bonferroni correction and pseudobulked “union” method. A minimum of 3 cells per gRNA-gene or element-gene pair was required for a discovery pair to pass quality control on each run.

### Comparisons between element-level measures

To aggregate SCEPTRE’s gRNA-gene results with FRACTEL, we used 10 million null simulations, a proportional bound of *k* = 0.3*n* and a minimum bound of 3. The intersection set of element-gene pairs across the FRACTEL results, Bonferroni test, and union test were used for all presented results; element-gene *p*-values were adjusted for multiple comparisons per test across this final intersection set using the Benjamini-Hochberg procedure. The standard error of *log*_2_ (*FC*) values across an element-gene pair was calculated across element-gene pairs containing more than 2 gRNA-gene pairs. A further set of results was generated with *k* = 0.5*n* to make Fig. S9.

## Declarations

### Availability of Data and Materials

The datasets used in this work were generated by the Impact of Genomic Variation on Function (IGVF) Consortium and are available through the IGVF Data Portal (https://data.igvf.org/). The specific datasets analyzed in this study correspond to IGVF accessions IGVFDS7132YVKO, IGVFDS0523KGZP, IGVFDS0790MEEM, IGVFDS3578UWZK, and IGVFDS3959LESA.

The FRACTEL implementation used for perturbation screen analyses is available at https://github.com/Gersbachlab-Bioinformatics/FRACTEL, and the code and data used in the simulation studies and power analyses are available at https://github.com/richardwdoty/fractel-simulations.

### Author contributions

A.S.A. conceived the research. A.S.A. and R.W.D. developed the FRACTEL method and statistical framework. M.A.t.W. developed the effect size estimator with supervision from A.S.A. R.W.D. implemented the power simulations, A.B. implemented the test, and M.A.t.W. implemented the effect size estimator, all with supervision from A.S.A. L.R.B. generated the experimental data with supervision from C.A.G. A.B. and L.R.B. processed the experimental data. M.A.t.W. ran and compared the statistical tests with assistance from A.B. and A.S.A. R.W.D. and M.A.t.W. interpreted the results with assistance from A.B. and A.S.A. R.W.D. wrote the manuscript with assistance from A.B. and M.A.t.W. under the supervision of A.S.A. A.S.A. supervised the project. All authors read and approved the final manuscript.

### Author information

**Authors and Affiliations**

**Department of Biostatistics & Bioinformatics, Duke University, Durham, 27705, NC, USA**

Richard W. Doty & Andrew S. Allen

**Center for Advanced Genomic Technologies, Duke University, Durham, 27705, NC, USA**

Richard W. Doty, Maria A. ter Weele, Alejandro Barrera, Lexi R. Bounds, Charles A. Gersbach & Andrew S. Allen

**Department of Biomedical Engineering, Duke University, Durham, 27705, NC, USA**

Maria A. ter Weele, Alejandro Barrera, Lexi R. Bounds & Charles A. Gersbach

**Xaira Therapeutics, South San Francisco, 94080, CA, USA**

Lexi R. Bounds

**Corresponding author**

Correspondence to Andrew S. Allen

## Supplementary material

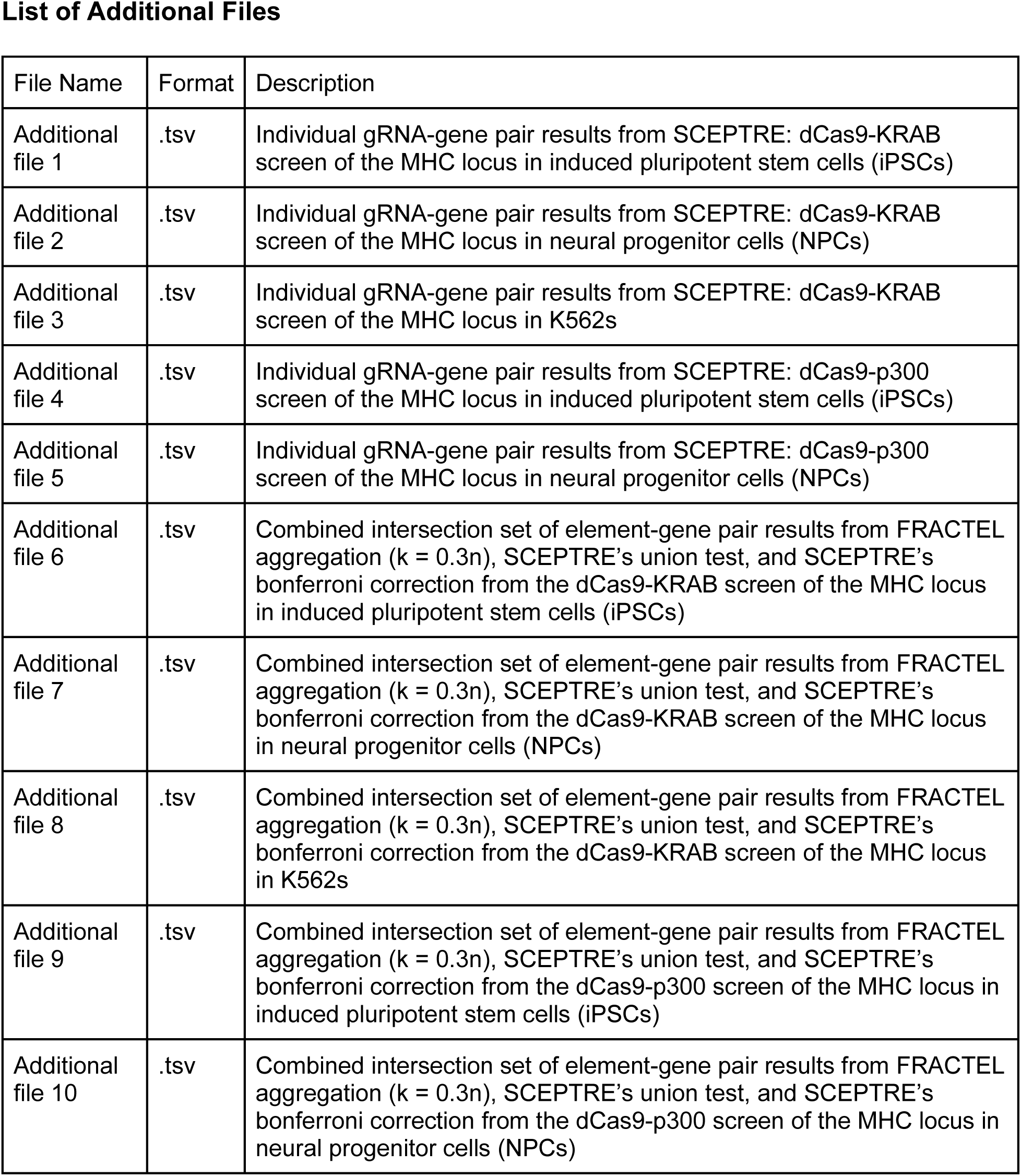
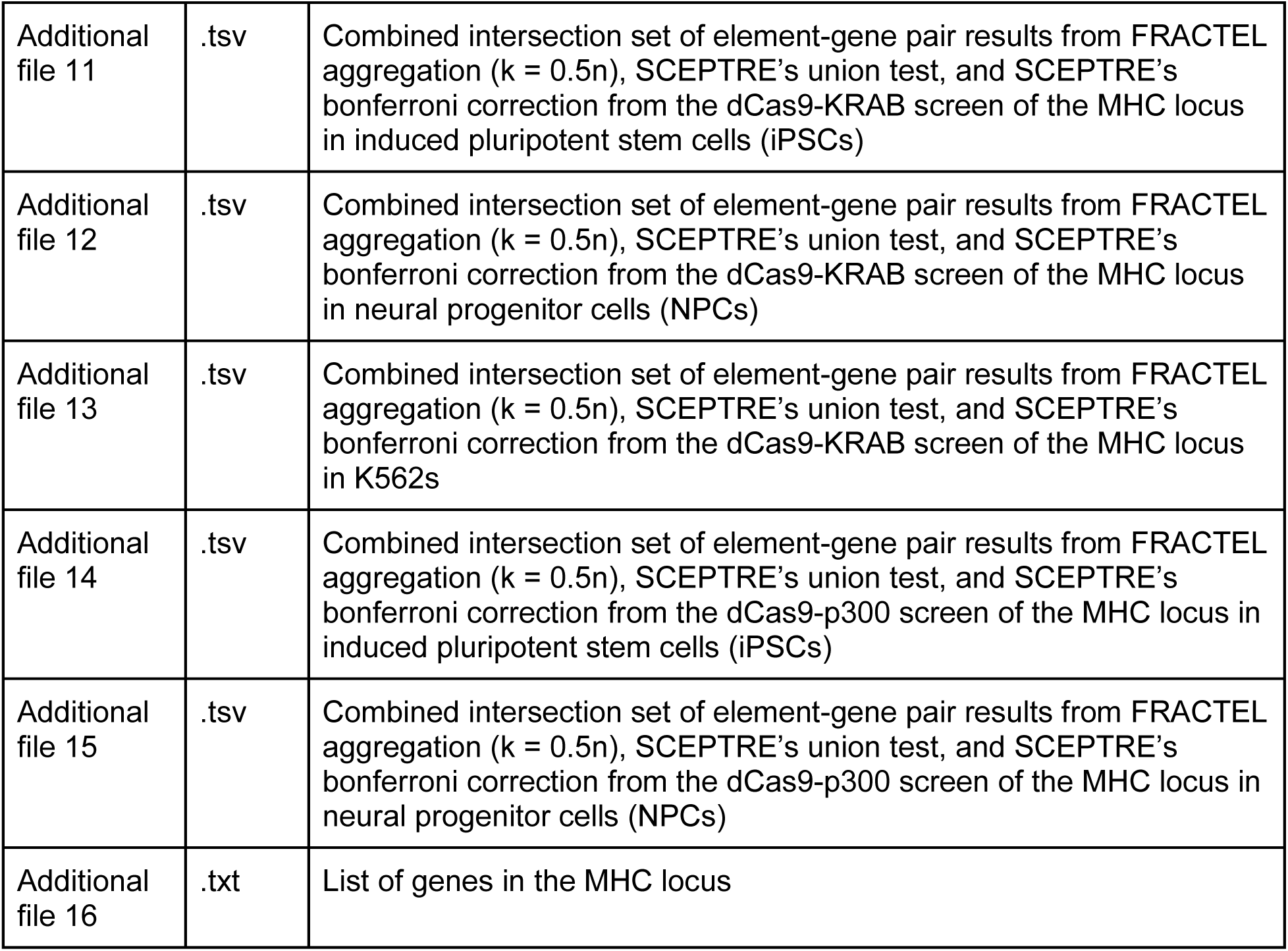

**List of Additional Files**
| File Name | Format | Description |
| --- | --- | --- |
| Additional file 1 | .tsv | Individual gRNA-gene pair results from SCEPTRE: dCas9-KRAB screen of the MHC locus in induced pluripotent stem cells (iPSCs) |
| Additional file 2 | .tsv | Individual gRNA-gene pair results from SCEPTRE: dCas9-KRAB screen of the MHC locus in neural progenitor cells (NPCs) |
| Additional file 3 | .tsv | Individual gRNA-gene pair results from SCEPTRE: dCas9-KRAB screen of the MHC locus in K562s |
| Additional file 4 | .tsv | Individual gRNA-gene pair results from SCEPTRE: dCas9-p300 screen of the MHC locus in induced pluripotent stem cells (iPSCs) |
| Additional file 5 | .tsv | Individual gRNA-gene pair results from SCEPTRE: dCas9-p300 screen of the MHC locus in neural progenitor cells (NPCs) |
| Additional file 6 | .tsv | Combined intersection set of element-gene pair results from FRACTEL aggregation ($k = 0.3n$), SCEPTRE's union test, and SCEPTRE's bonferroni correction from the dCas9-KRAB screen of the MHC locus in induced pluripotent stem cells (iPSCs) |
| Additional file 7 | .tsv | Combined intersection set of element-gene pair results from FRACTEL aggregation ($k = 0.3n$), SCEPTRE's union test, and SCEPTRE's bonferroni correction from the dCas9-KRAB screen of the MHC locus in neural progenitor cells (NPCs) |
| Additional file 8 | .tsv | Combined intersection set of element-gene pair results from FRACTEL aggregation ($k = 0.3n$), SCEPTRE's union test, and SCEPTRE's bonferroni correction from the dCas9-KRAB screen of the MHC locus in K562s |
| Additional file 9 | .tsv | Combined intersection set of element-gene pair results from FRACTEL aggregation ($k = 0.3n$), SCEPTRE's union test, and SCEPTRE's bonferroni correction from the dCas9-p300 screen of the MHC locus in induced pluripotent stem cells (iPSCs) |
| Additional file 10 | .tsv | Combined intersection set of element-gene pair results from FRACTEL aggregation ($k = 0.3n$), SCEPTRE's union test, and SCEPTRE's bonferroni correction from the dCas9-p300 screen of the MHC locus in neural progenitor cells (NPCs) |
| Additional file 11 | .tsv | Combined intersection set of element-gene pair results from FRACTEL aggregation ($k = 0.5n$), SCEPTRE's union test, and SCEPTRE's bonferroni correction from the dCas9-KRAB screen of the MHC locus in induced pluripotent stem cells (iPSCs) |
| Additional file 12 | .tsv | Combined intersection set of element-gene pair results from FRACTEL aggregation ($k = 0.5n$), SCEPTRE's union test, and SCEPTRE's bonferroni correction from the dCas9-KRAB screen of the MHC locus in neural progenitor cells (NPCs) |
| Additional file 13 | .tsv | Combined intersection set of element-gene pair results from FRACTEL aggregation ($k = 0.5n$), SCEPTRE's union test, and SCEPTRE's bonferroni correction from the dCas9-KRAB screen of the MHC locus in K562s |
| Additional file 14 | .tsv | Combined intersection set of element-gene pair results from FRACTEL aggregation ($k = 0.5n$), SCEPTRE's union test, and SCEPTRE's bonferroni correction from the dCas9-p300 screen of the MHC locus in induced pluripotent stem cells (iPSCs) |
| Additional file 15 | .tsv | Combined intersection set of element-gene pair results from FRACTEL aggregation ($k = 0.5n$), SCEPTRE's union test, and SCEPTRE's bonferroni correction from the dCas9-p300 screen of the MHC locus in neural progenitor cells (NPCs) |
| Additional file 16 | .txt | List of genes in the MHC locus |

## Supplementary Tables

**Table S1:** Empirical type I error rates across baseline expression levels for varying values of n. Values are estimated from simulations under the null with *α* = 0.05.

| n | Baseline expression (UMI) |  |  |
| --- | --- | --- | --- |
|  | Low | Moderate | High |
| 5 | 0.0500 | 0.0496 | 0.0501 |
| 6 | 0.0496 | 0.0521 | 0.0497 |
| 7 | 0.0495 | 0.0493 | 0.0499 |
| 8 | 0.0491 | 0.0498 | 0.0482 |
| 9 | 0.0486 | 0.0496 | 0.0498 |
| 10 | 0.0497 | 0.0516 | 0.0513 |
| 11 | 0.0493 | 0.0502 | 0.0494 |
| 12 | 0.0502 | 0.0487 | 0.0505 |
| 13 | 0.0489 | 0.0506 | 0.0498 |
| 14 | 0.0512 | 0.0505 | 0.0498 |
| 15 | 0.0514 | 0.0516 | 0.0510 |
| 16 | 0.0504 | 0.0506 | 0.0490 |
| 17 | 0.0513 | 0.0499 | 0.0493 |
| 18 | 0.0489 | 0.0505 | 0.0488 |
| 19 | 0.0497 | 0.0510 | 0.0495 |
| 20 | 0.0494 | 0.0496 | 0.0509 |

**Figure S1:**
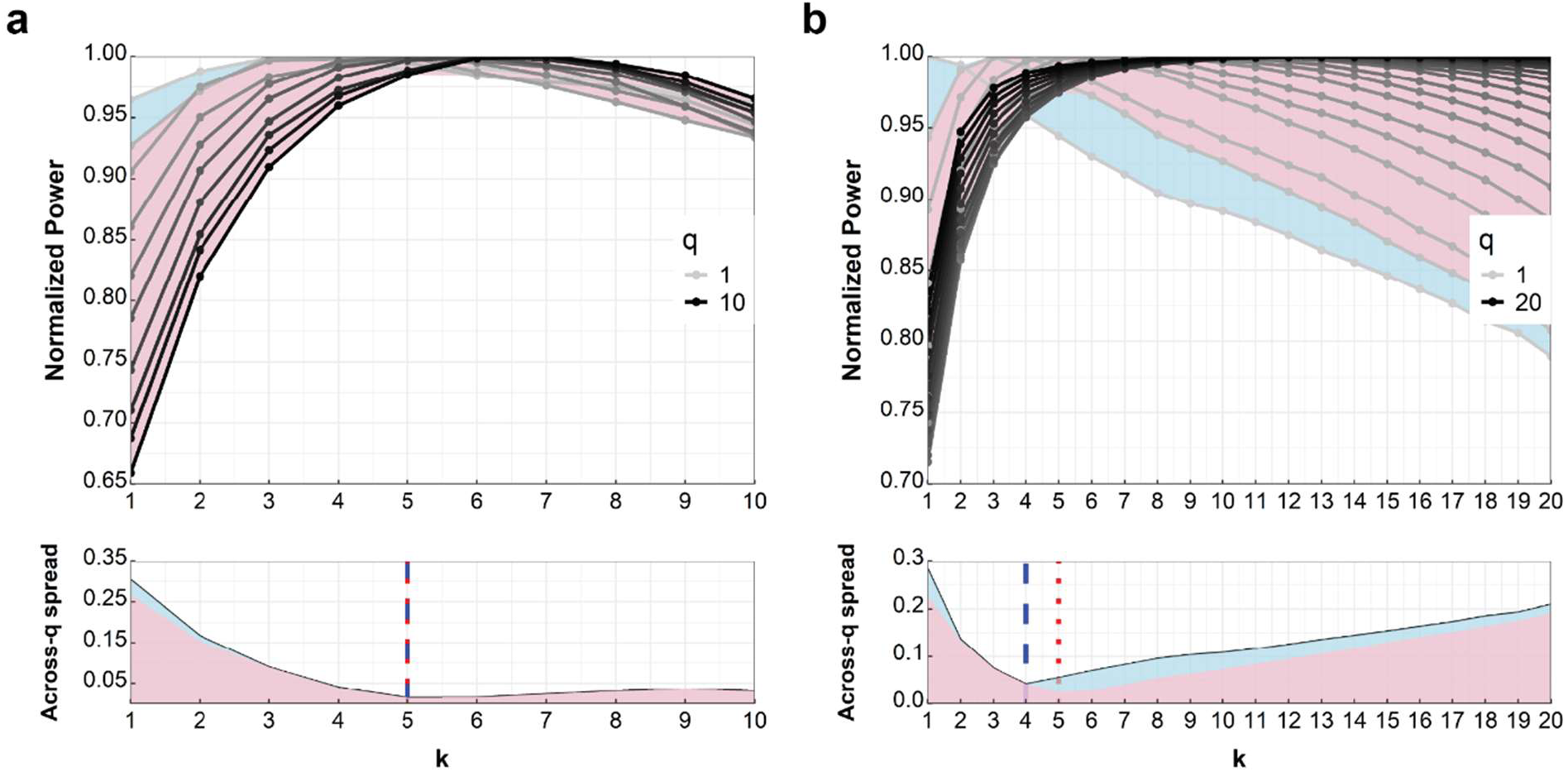
Sensitivity of the optimal bound parameter k. Normalized power curves are shown as in Fig. 2b, with each line scaled by its maximum across k for a fixed number of active gRNAs (q). The shaded region represents the range of normalized power across architectures, and the lower subplot shows the width of this range (max–min) as a function of k. (**a)** Reduced baseline expression. In this setting, the minimum spread occurs near k = 3–4, similar to Fig. 2b. Excluding the single active gRNA case (q = 1) does not materially shift the location of this minimum. However, the spread remains relatively flat for larger k, indicating reduced sensitivity of this criterion for selecting a single preferred value of k under lower baseline expression. **(b)** Increased guide count (n = 20). The minimum spread occurs near k = 4–5, with the precise location depending on whether the q = 1 case is included. In contrast to the n = 10 setting, the spread increases more rapidly for larger k, indicating stronger penalization of higher k values when signal is sparse. These results suggest that the choice of k becomes more sensitive to both guide count and assumptions about sparsity as n increases.

**Figure S2:**
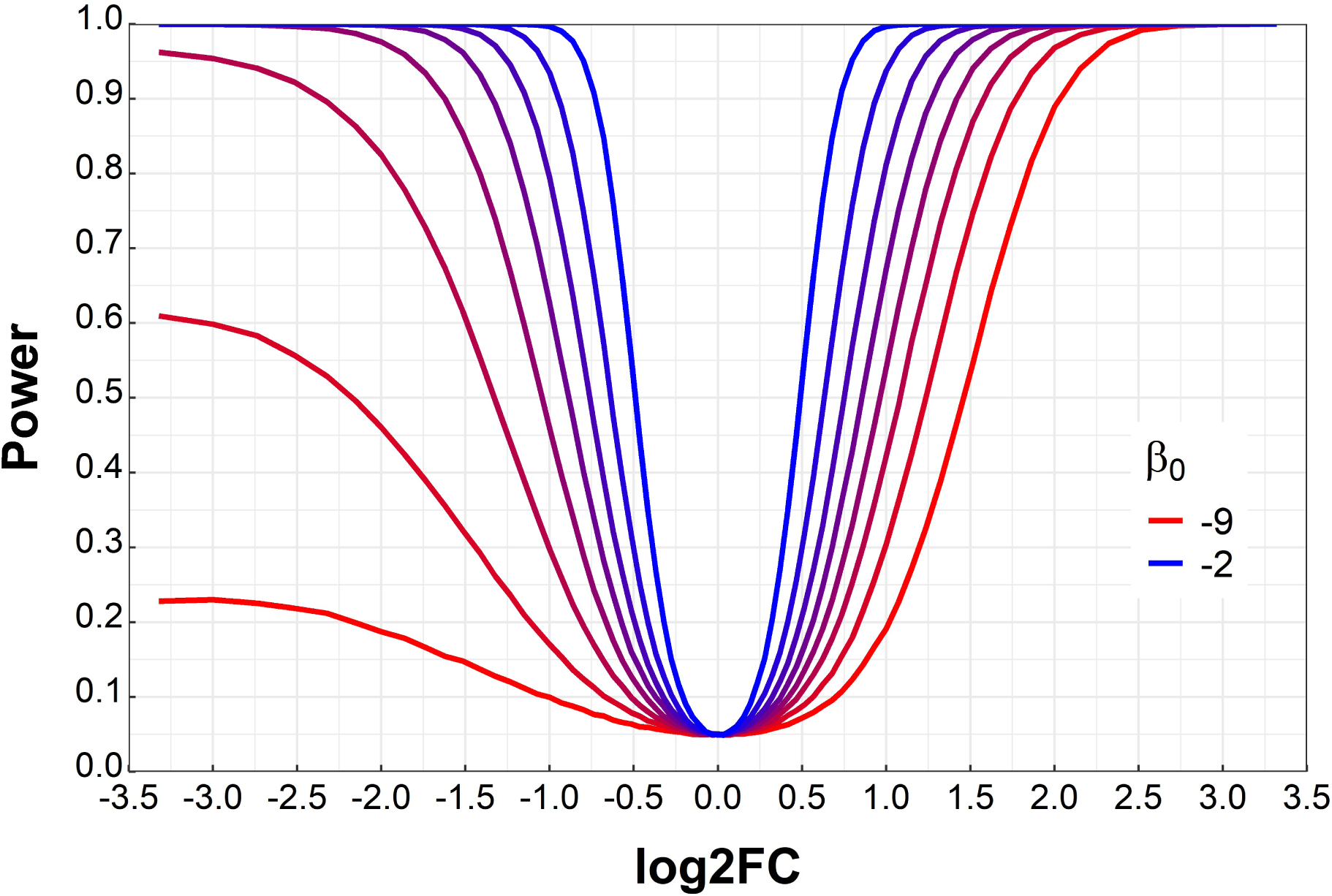
Power curves across different baseline expressions. All curves were simulated with identical parameters except *β*_0_, which ranges from UMI≈0.04 to 40.6. For low values of *β*_0_repression effects (*log*_2_ (*FC*) < 0) show clear asymmetry compared to activation effects (*log*_2_ (*FC*) > 0).This asymmetry disappears as *β*_0_ increases.

**Figure S3:**
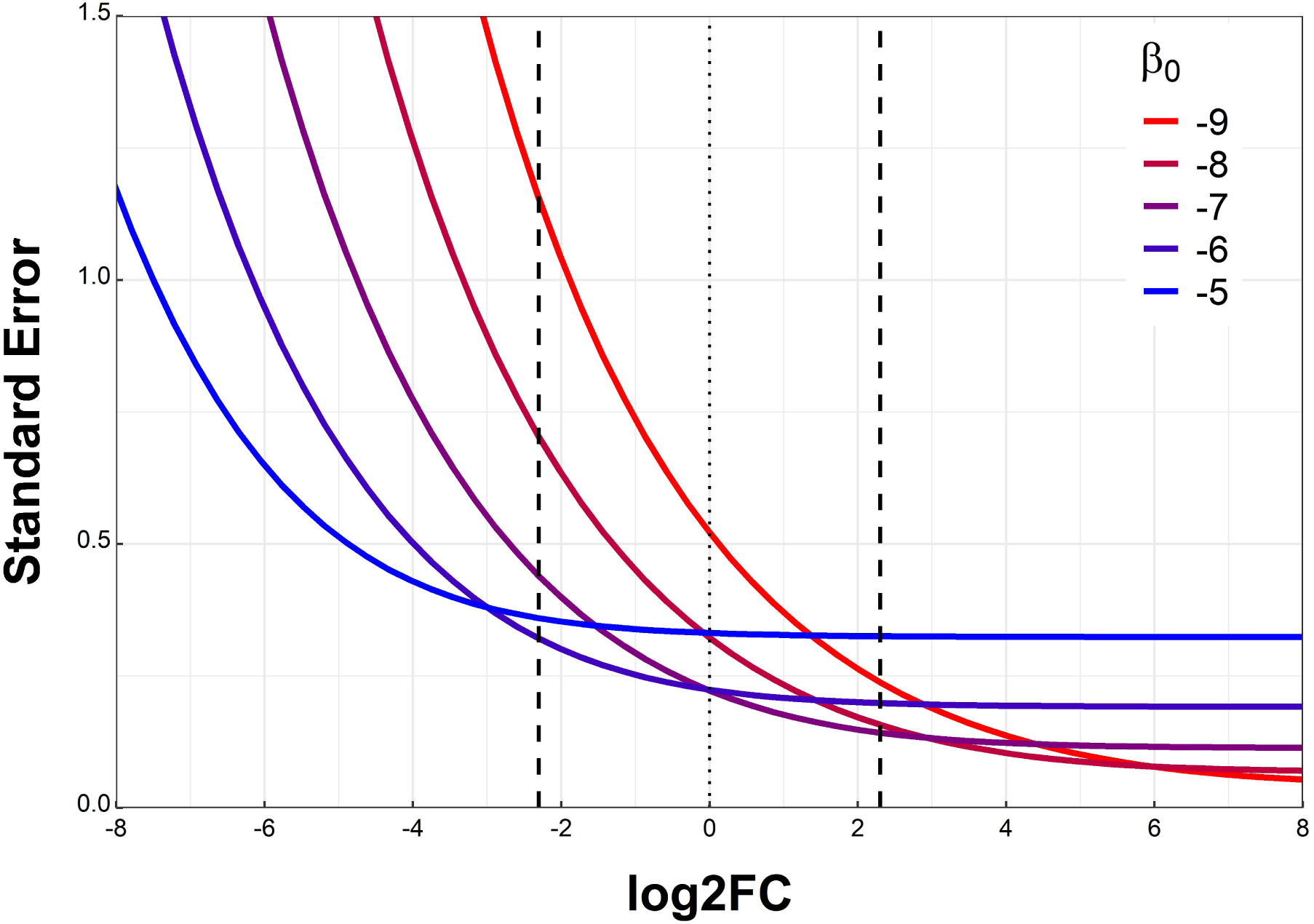
Standard error for varying baseline expressions. While each curve corresponding to a different baseline expression shows that standard error increases as log2FC decreases and plateaus as it increases, the effect becomes more visible in practical effect ranges for lower baseline expression. The vertical black lines correspond to FC=0.1 and FC=10. For all effects on *FC* ∈ (0.1,10), the standard error is effectively constant for UMI >= 2.

**Figure S4:**
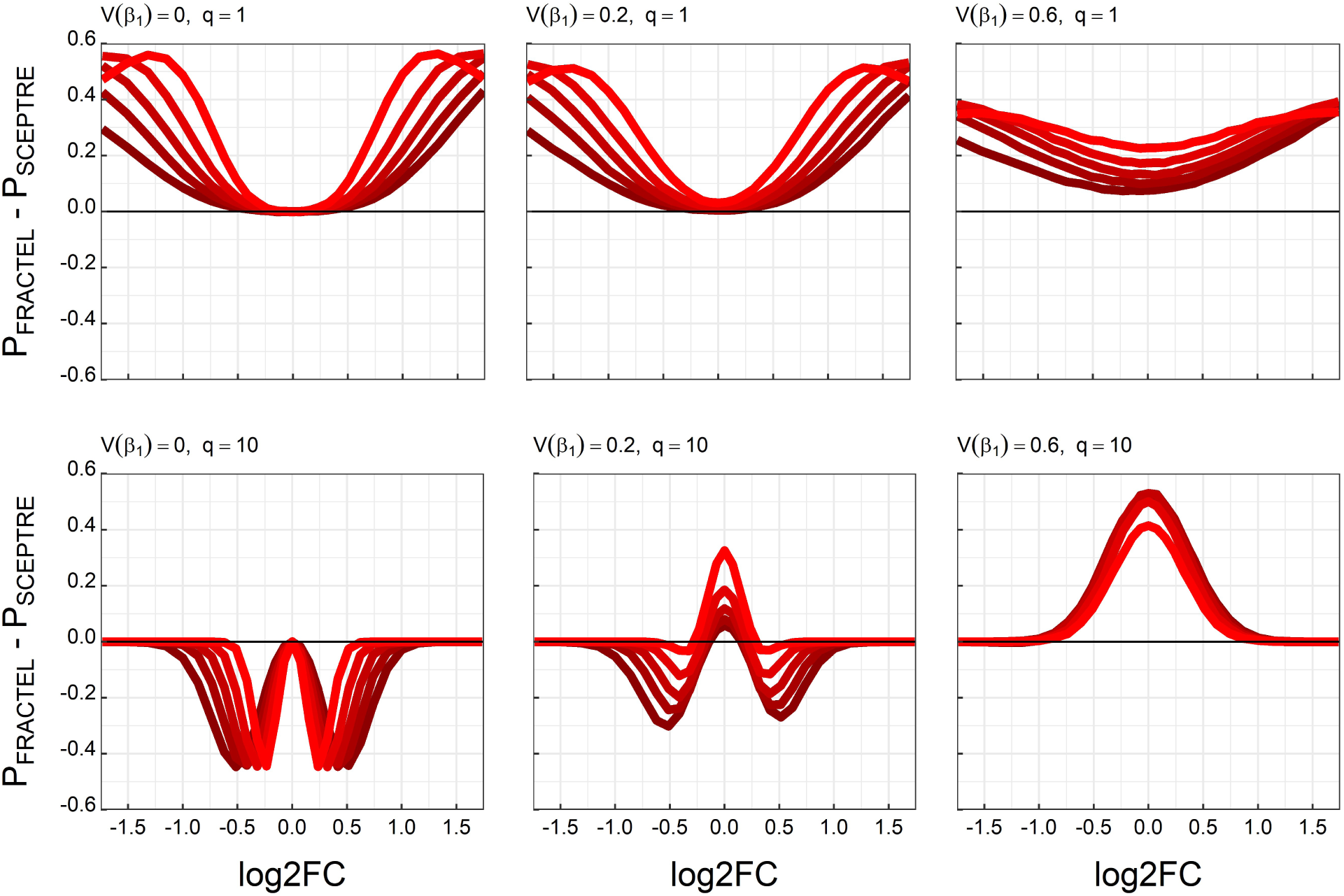
Difference in power between FRACTEL and union-based aggregation for n = 10. Curves show *ΔPower* = *P*_FRACTEL_ − *P*_SCEPTRE_across log2 fold-change effect sizes with brighter colors reflecting higher baseline expressions. The left column corresponds to sparse signal (q = 1), and the right column corresponds to dense signal (q = 10). Rows vary the heterogeneity of gRNA effect sizes. The top row shows no variance in effect size (all active guides share the same effect). The middle row introduces moderate heterogeneity, with effect sizes drawn on the log2FC scale from a Normal(*β*_l_, 0.2) distribution. The bottom row shows high heterogeneity, with effect sizes drawn on the log2FC scale from a Normal(*β*_l_, 0.6). When effects are homogeneous and many guides are active (top right), the union method is more powerful across most effect sizes, reflecting efficient aggregation of consistent signal. In contrast, FRACTEL outperforms the union method in sparse settings (q = 1; left column) and as heterogeneity increases (middle and bottom rows), where variation in effect direction or magnitude reduces the effectiveness of union-based aggregation. Under high variance (bottom row), FRACTEL maintains higher power across nearly all effect sizes, particularly when multiple guides are active.

**Figure S5:**
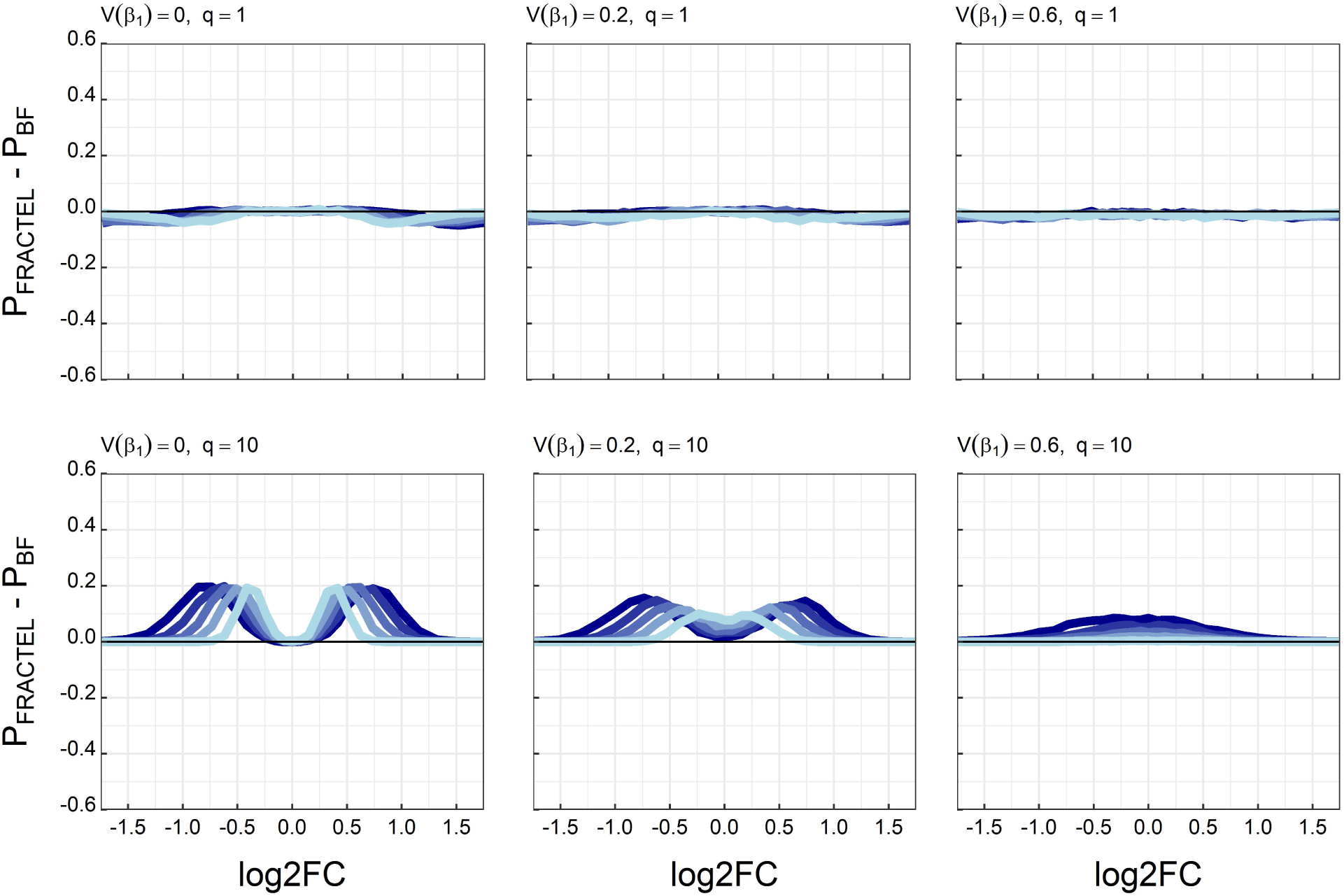
Difference in power between FRACTEL and Bonferroni-corrected minimum p-value aggregation for n = 10. Curves show *ΔPower* = *P*_FRACTEL_ − *P*_BF_across log2 fold-change effect sizes with brighter colors reflecting higher baseline expressions. The top row corresponds to sparse signal (q = 1), and the bottom row corresponds to dense signal (q = 10). Columns vary the heterogeneity of gRNA effect sizes. The left column shows no variance in effect size, the middle column introduces moderate heterogeneity, and the right column shows high heterogeneity. For sparse signal (q = 1), FRACTEL and Bonferroni aggregation perform similarly across all levels of effect size variability. For dense signal (q = 10), FRACTEL achieves higher power, reflecting its ability to aggregate signal across multiple active guides. However, this advantage diminishes as effect size variability increases.

**Figure S6.**
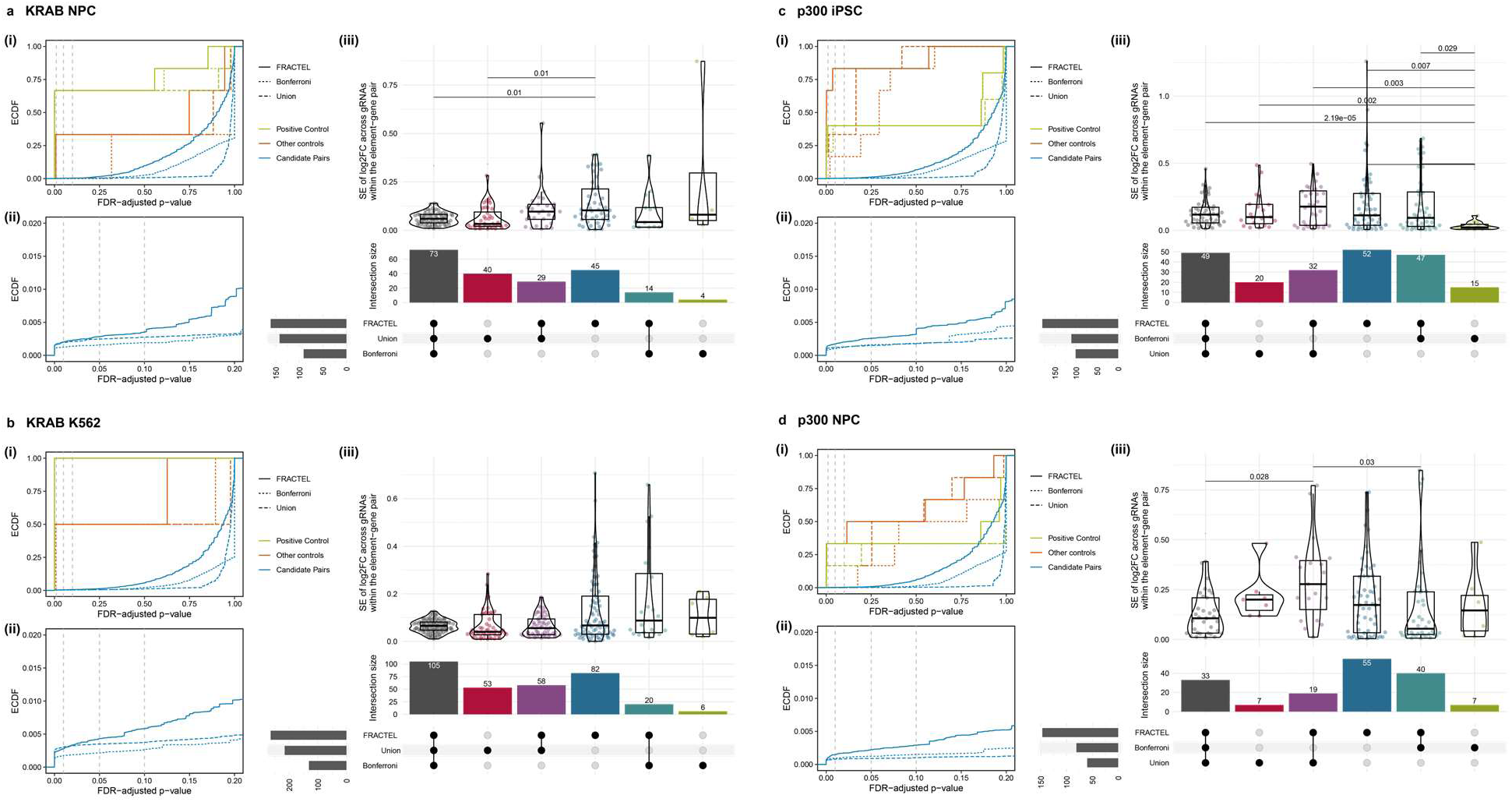
Comparison of aggregation methods across 4 additional screens. Perturb-seq screens of the MHC locus with dCas9-KRAB in **(a)** neural progenitor cells and **(b)** K562s, and with dCas9-p300 in **(c)** induced pluripotent stem cell (iPSCs) and **(d)** neural progenitor cells (NPCs)^1^**. (i)** Empirical cumulative distribution function (ECDF) of the three aggregation methods across FDRs. **(ii)** Plot of ECDF in **(i)** for non-control elements with FDR < 0.2. **(iii)** Comparison of standard error of the effect sizes (*log*_2_ (*FC*)) between gRNAs within an element-gene pair, for exclusive intersection sets of significant (FDR < 0.05) elements called by each of the three methods, along with an upset plot to show set sizes. Significant pairwise Mann-Whitney U-tests with a Holm correction are displayed (*p*_adj_ < 0.05). Not shown is the exclusive intersection of the union and Bonferroni methods (**(a)** n = 0, **(b)** n = 0, **(c)** n = 0, **(d)** n = 1).

**Figure S7.**
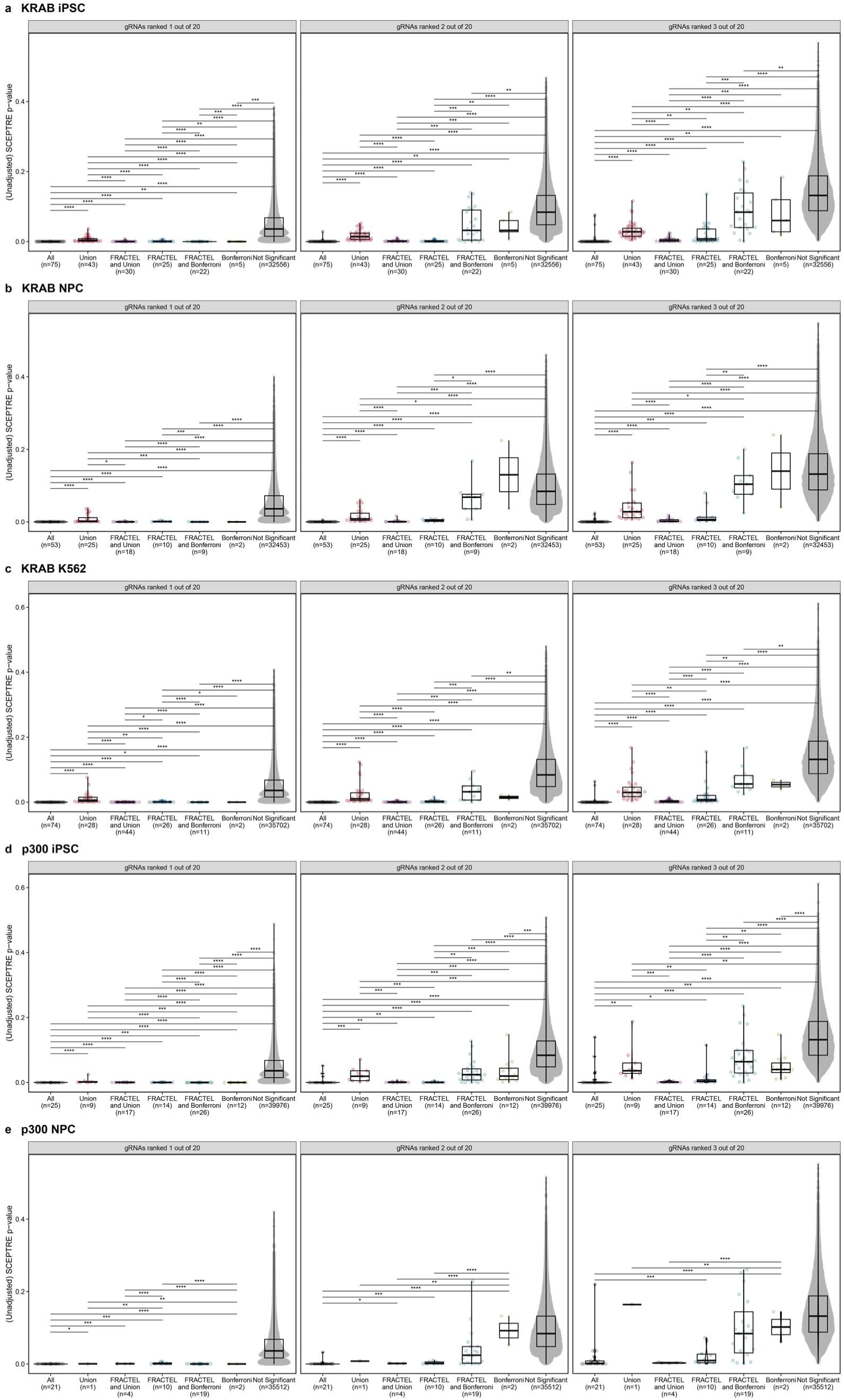
Comparison of aggregation methods’ ordered p-values. The first three ordered SCEPTRE gRNA-gene p-values from Perturb-seq screens of the MHC locus with dCas9-KRAB in **(a)** induced pluripotent stem cells (iPSCs) **(b)** neural progenitor cells (NPCs), and **(c)** K562s, and with dCas9-p300 in **(c)** iPSCs and **(d)** NPCs^1^. Data were analyzed with SCEPTRE (see Methods) and aggregated to element-gene statistics using either SCEPTRE’s Bonferroni method, SCEPTRE’s pseudobulked union method, or FRACTEL (k = 0.3n). Only element-gene pairs with 20 complete gRNA-gene pairs are shown. Pairwise Mann-Whitney U-tests were used to compare p-values between exclusive sets called by each of the methods at FDR = 0.05 within each order (significances: **** p< 0.0001, *** p < 0.001, ** p < 0.01, * p < 0.05 after Holm adjustment). Not shown is the exclusive intersection of the union and Bonferroni methods (**(a)** n = 1, **(b)** n = 0, **(c)** n = 0, **(d)** n = 0, **(e)** n = 0).

**Figure S8:**
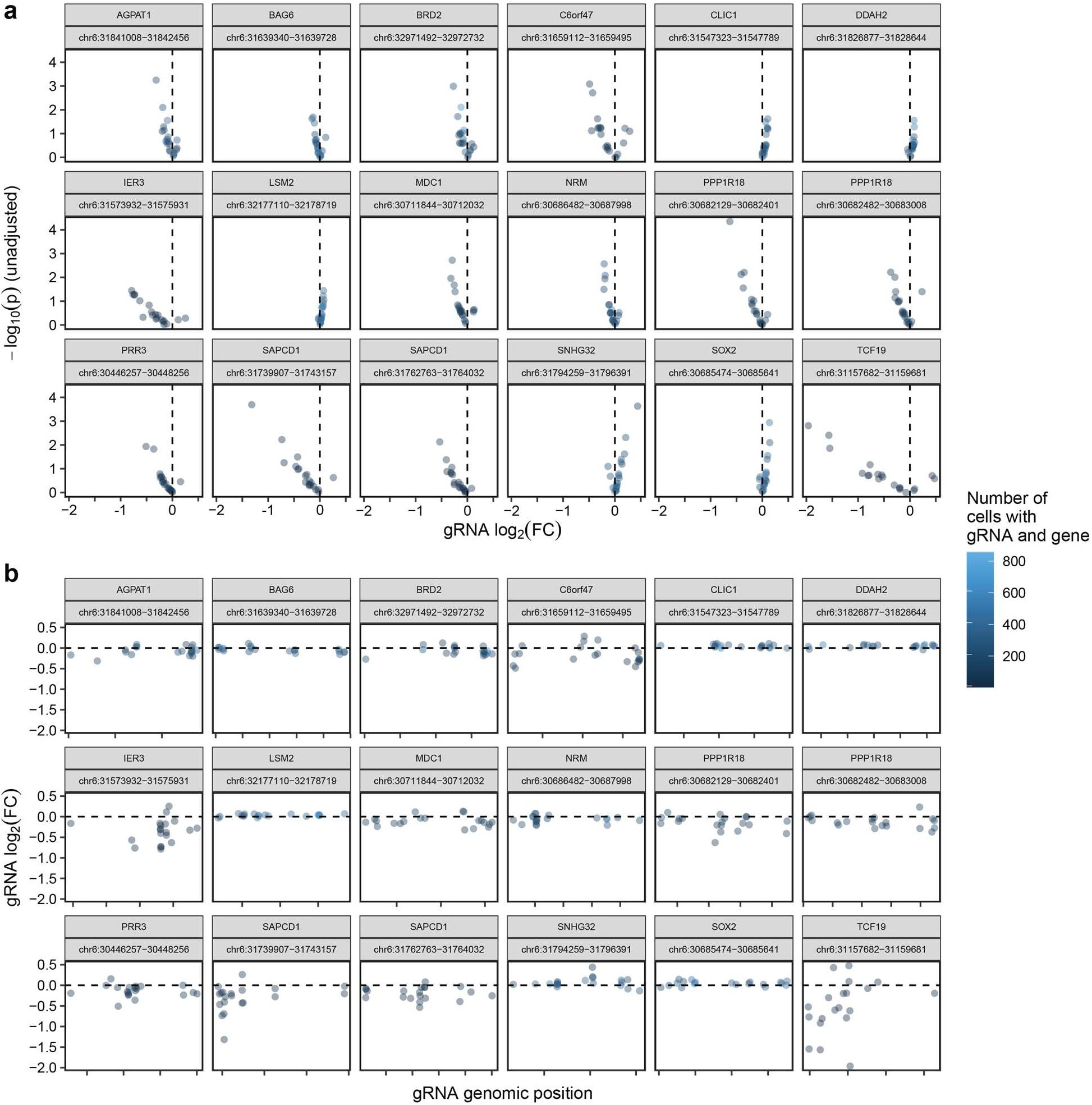
Elements uniquely significant in the union analysis. Examples of some element-gene pairs significant in the union analysis (FDR < 0.05) but not FRACTEL (FDR > 0.2) in a dCas9-KRAB Perturb-seq screen of the MHC locus in neural progenitor cells. Points represent gRNA-gene pairs, colored by the number of cells containing the gRNA and gene, and grouped by element-gene pair. Element-gene pairs are labeled with the affected gene symbol and coordinates of the targeted genomic element. **(a)** Volcano plots showing the unadjusted significances (-*log*_l0_ *p*) and effect sizes (*log*_2_ (*FC*)) from SCEPTRE for individual gRNA-gene pairs. **(b)** Plots of SCEPTRE gRNA-gene effect sizes (*log*_2_ (*FC*)), displayed by genomic position across the element within element-gene pairs.

**Figure S9:**
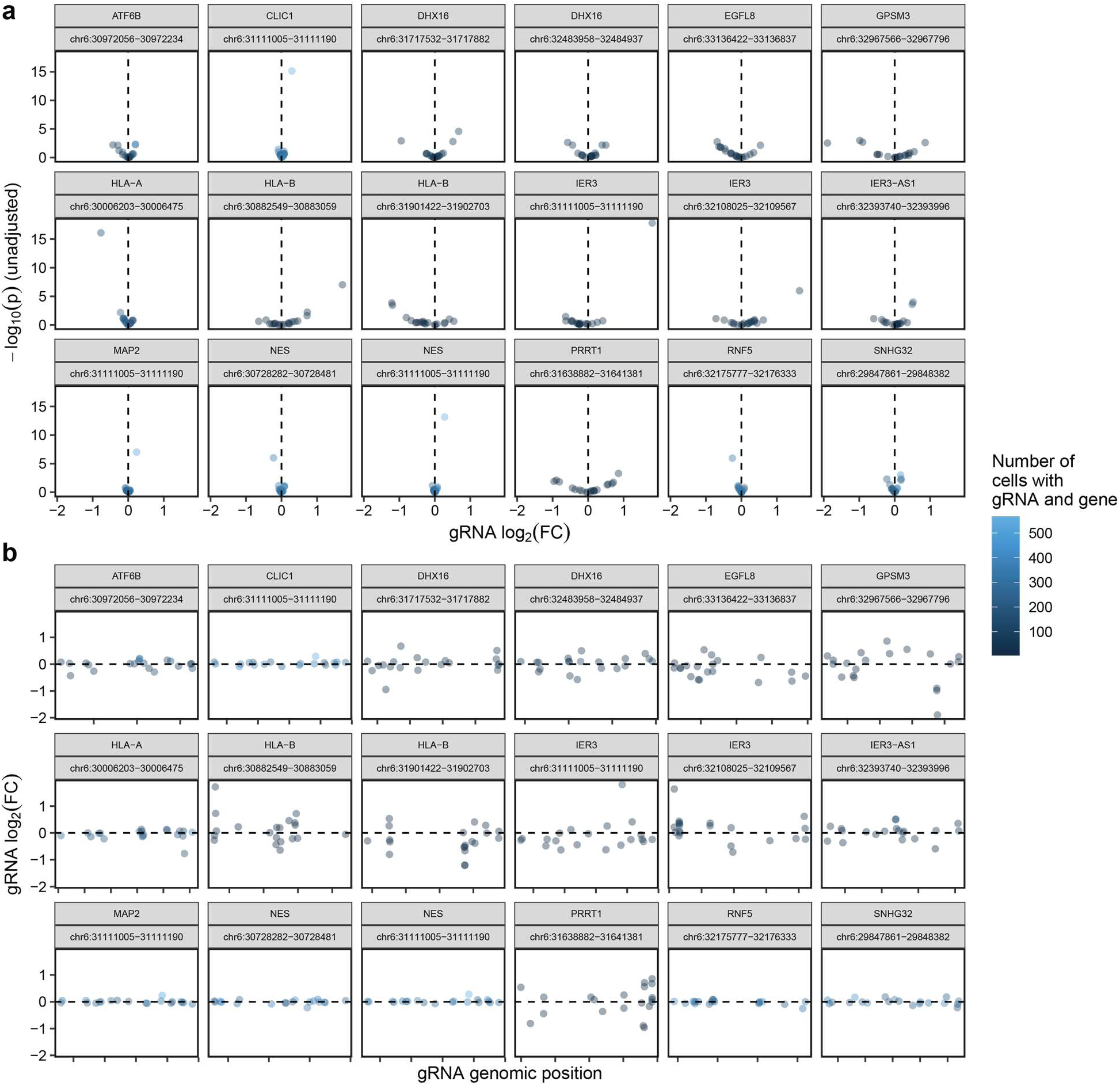
Elements uniquely significant in FRACTEL. Examples of some element-gene pairs significant in FRACTEL aggregation (FDR < 0.05) but not the union analysis (FDR > 0.2) in a dCas9-KRAB Perturb-seq screen of the MHC locus in neural progenitor cells. Points represent gRNA-gene pairs, colored by the number of cells containing the gRNA and gene, and grouped by element-gene pair. Element-gene pairs are labeled with the affected gene symbol and coordinates of the targeted genomic element. **(a)** Volcano plots showing the unadjusted significances (-*log*_l0_ *p*) and effect sizes (*log*_2_ (*FC*)) from SCEPTRE for individual gRNA-gene pairs. **(b)** Plots of SCEPTRE gRNA-gene effect sizes (*log*_2_ (*FC*)), displayed by genomic position across the element within element-gene pairs.

**Figure S10:**
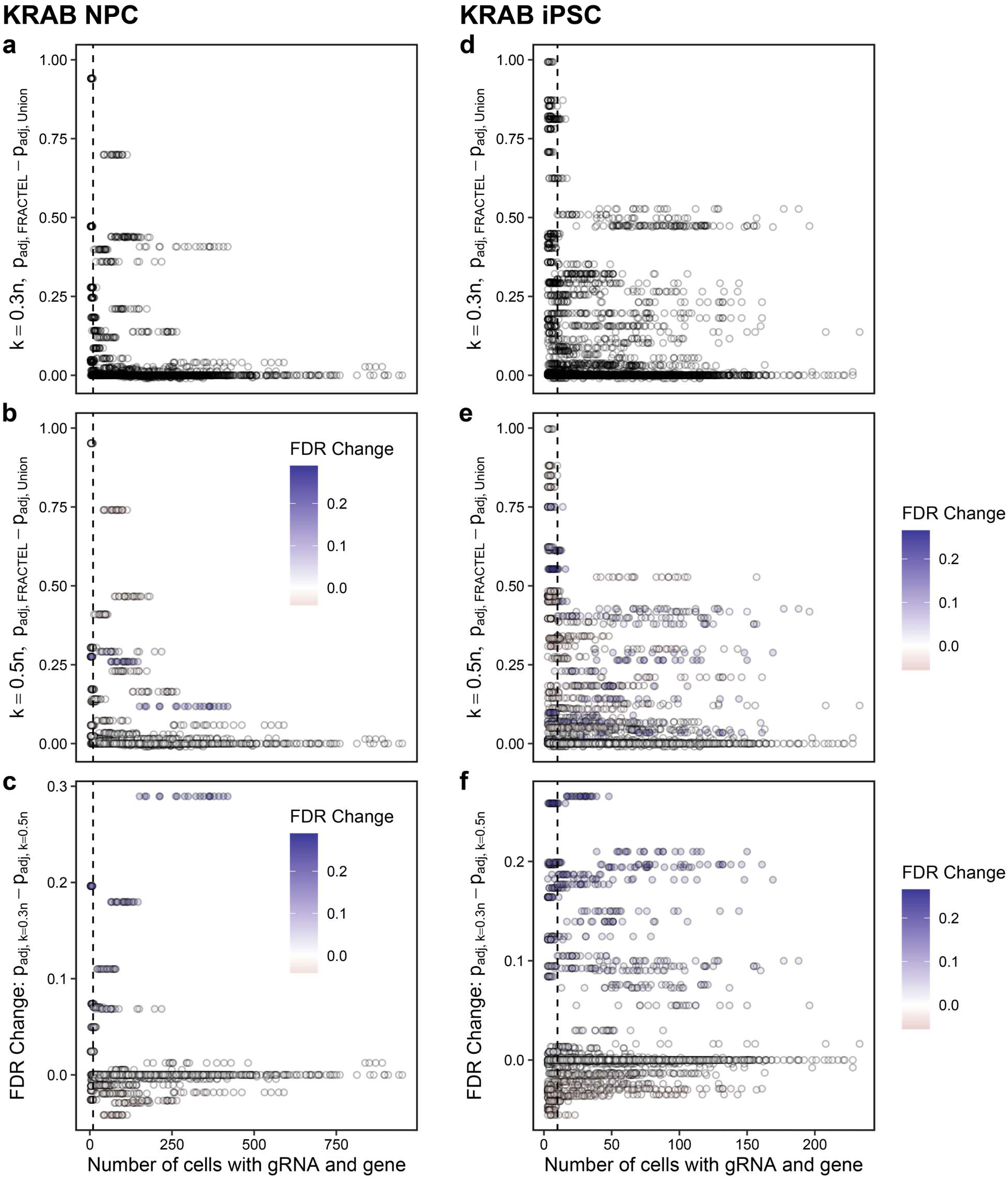
Effect of tunable bound on highly significant union elements. Plotted are element-gene pairs with FDR_Union_ < 0.01 from the dCas9-KRAB Perturb-seq screen of the MHC locus in **(a-c)** neural progenitor cells (NPCs) and **(d-f)** induced pluripotent stem cells (iPSCs). Plotted is the number of cells with the gRNA and gene in the gRNA-level SCEPTRE analysis and **(a,f)** the difference between FRACTEL and union FDRs with *k* = 0.3*n*, **(b,e)** the difference between FRACTEL and union FDRs with *k* = 0.3*n*, colored by the FDR change between *k* = 0.5*n* and *k* = 0.3*n*, and **(c,f)** the FDR change between *k* = 0.5*n* and *k* = 0.3*n*, colored identically.

**Figure S11:**
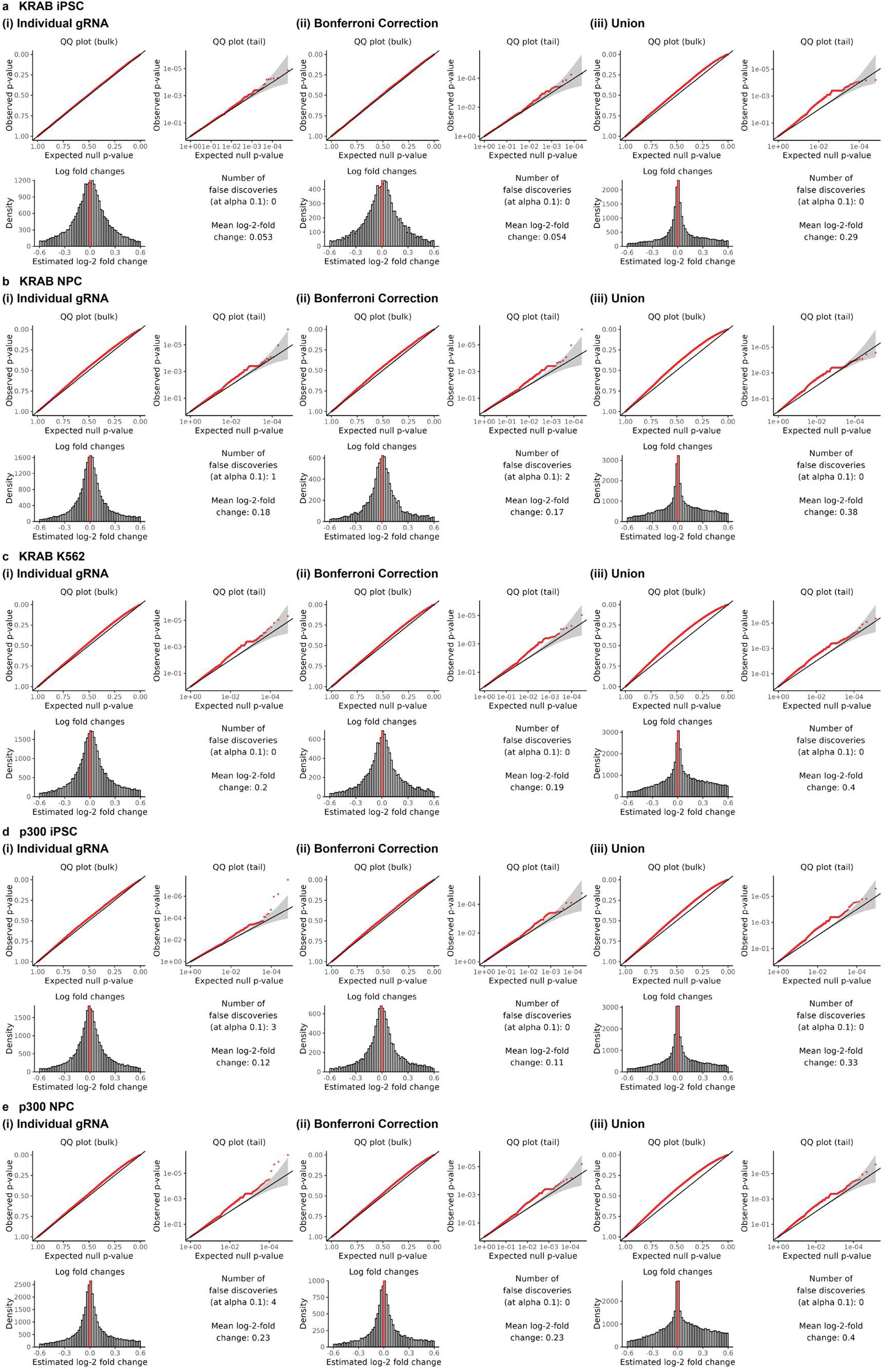
Calibration checks for SCEPTRE analysis. Calibration checks from SCEPTRE for each of the 5 screens Perturb-seq screens of the MHC locus with dCas9-KRAB in **(a)** induced pluripotent stem cells (iPSCs) **(b)** neural progenitor cells (NPCs), and **(c)** K562s, and with dCas9-p300 in **(c)** iPSCs and **(d)** NPCs^48^.

